# A geostatistical approach to modelling human Holocene migrations in Europe using ancient DNA

**DOI:** 10.1101/826149

**Authors:** Fernando Racimo, Jessie Woodbridge, Ralph M. Fyfe, Martin Sikora, Karl-Göran Sjögren, Kristian Kristiansen, Marc Vander Linden

## Abstract

The European continent was subject to two major migrations of peoples during the Holocene: the northwestward movement of Anatolian farmer populations during the Neolithic and the westward movement of Yamnaya steppe peoples during the Bronze Age. These movements changed the genetic composition of the continent’s inhabitants, via admixture and population replacement processes. The Holocene was also characterized by major changes in vegetation composition, which altered the environment occupied by the original hunter-gatherer populations. Here, we use a combination of paleogenomics and geostatistical modelling to produce detailed maps of the movement of these populations over time and space, and to understand how these movements impacted the European vegetation landscape. We find that the dilution of hunter-gatherer ancestries and the Yamnaya steppe migration coincided with a reduction in the amount of broad-leaf forest and an increase in the amount of pasture lands in the continent. We also show that climate played a role in these vegetational changes. Additionally, we find that the spread of Neolithic farmer ancestry had a two-pronged wavefront, in agreement with similar findings based on patterns of the cultural spread of farming from radiocarbon-dated archaeological sites. With thousands of ancient genomes publicly available and in production, we foresee that the integration of ancient DNA with geostatistical techniques and large-scale archaeological datasets will revolutionize the study of ancient populations movements, and their effects on local fauna and flora.

## Introduction

Recent paleogenomic studies - often involving hundreds of individuals - have provided new insights into the nature of large-scale population movements during the Holocene (Allentoft et al., 2015; Haak et al., 2015; Mathieson, Lazaridis, et al., 2015; de Barros Damgaard et al., 2018; Mathieson, Alpaslan-Roodenberg, et al., 2018; de Barros Damgaard et al., 2018). Although we cannot directly observe migrations of people, as they occurred in the past, we can observe spatial changes in genomic ancestries over time. Under a latent mixed-membership model, these ancestries represent inferred, unobserved ancestral groups that contributed different amounts of genetic material to each of the individuals analyzed, and are found by fitting allele frequency patterns to the genomes of these individuals. Changes in ancestry can indicate when people from distant lands started producing offspring in new, distant locations from their previous homelands. Nevertheless, considerable care must be taken while interpreting these inferred ancestries as true ancestral populations (Daniel J Lawson, Van Dorp, and Falush, 2018).

Analyses based on patterns of ancestry as well as various other sources of population genetic evidence based on differential allele sharing indicate a wave of populations from the Middle East entered Europe starting ~8,500 years ago (Sikora et al., 2014; Lazaridis, Patterson, et al., 2014). Furthermore, studies based on large-scale datasets of radiocarbon-dated domestic plants, animals and finds from associated contexts (Vander Linden and Silva, 2018) indicate that this migration wave spread farming practices into the region, initiating the Neolithic revolution in Europe (Ammerman and Cavalli-Sforza, 1971). A second massive wave of movement occurred at the beginning of the Bronze Age, when populations associated with the Yamnaya culture in the Pontic steppe entered the continent from the east (Anthony, 2010; Shishlina, 2008; Kristiansen and Larsson, 2005). These groups might have introduced horse-herding and proto-Indo-European languages as they moved westward, and are associated with the Corded Ware culture in central and northern Europe and, later on, the Bell Beaker complex in northwestern Europe (Haak et al., 2015; Allentoft et al., 2015; Olalde et al., 2018; Vandkilde, 2007).

In parallel, over the last 10,000 years, the European continent has undergone major changes in its land-cover composition. Recent pollen-based studies suggest a dramatic reduction of broad-leaf forests in Europe occurred from about 6,000 BP until the present (N. Roberts et al., 2018). The deforestation intensified at around 2,200 BP, resulting in a replacement of these forests by pasture and arable land throughout the continent (Marquer et al., 2017; R. M. Fyfe, Woodbridge, and N. Roberts, 2015). This process, however, did not occur at the same rate throughout all regions of the continent. For example, while considerable decreases in broad-leaf forests occurred in central Europe starting around 4,000 BP, the Atlantic seaboard was predominantly occupied by semi-open vegetation since well before this occurred, while southern Scandinavia underwent much less considerable reductions in forest cover, at least until the Middle Ages (R. M. Fyfe, Woodbridge, and N. Roberts, 2015; Anne Birgitte Nielsen et al., 2012; R. M. Fyfe, Twiddle, et al., 2013). Presumably, this phenomenon was partly effected by new human land use activities involving forest clearance and the establishment of farming and herding practices. Most of these activities were introduced into the continent via migration of foreign populations: up until 8,000 BP, Europe was largely populated by groups of hunter-gatherers living at relatively low densities, with likely limited effects on their surrounding flora and fauna (although see Bishop, Church, and Rowley-Conwy (2015) and Warren et al. (2014)). Additionally, changes in climate patterns may have also played a role. Until now, however, few efforts have been carried out to explicitly link changes in paleo-vegetation to particular human population movements, or to distinguish between climatic and human-based factors (but see Marquer et al. (2017)).

Paleogenomic ancestry inferences provide a way to trace these migrations in unprecedented spatial detail. In this study, we aim to model the major Holocene migrations using geostatistical methods - like spatiotemporal kriging and hierarchical Bayesian modeling - which are commonly used to model environmental processes. We use these methods to better study these migrations, their relationship to the spread of farming practices and to Holocene vegetation change. Additionally, we estimate their front speed, to better understand how these may have occurred, and compare our results to reconstructions of cultural dispersal obtained from radiocarbon-dated archaeological sites.

Our modeling approach suggests that the decline in broad-leaf forest was concurrent with a decline in hunter-gatherer ancestry, perhaps as a consequence of outside migrations, starting in the Neolithic. The major conversion from forest to pasture land, however, did not occur until fairly late in the Holocene, about 4,000 years ago. Our results suggest that it may have been the fast movement of steppe peoples that kick-started this major conversion. Natural variations in climate patterns during this period also played a role, however. For example, increases in temperatures seem to be related to increases in scrubland, pasture and arable land. Although our analyses do not include other important variables that might have played important roles - like increases in population density in the last 2,000 years - they do point to important factors that may have affected land-cover change in the past 10,000 years and pave the way for future geostatistical studies integrating paleogenomics with other paleo-disciplines.

## Results

We downloaded publicly available ancient and present-day genomic sequences typed at the Human Origins SNP array (Mathieson, Lazaridis, et al., 2015; Lazaridis, Patterson, et al., 2014; Patterson et al., 2012; Allentoft et al., 2015; Haak et al., 2015; Mathieson, Alpaslan-Roodenberg, et al., 2018; Olalde et al., 2018; Lazaridis, Nadel, et al., 2016). We performed unsupervised latent ancestry estimation on each of these sequences using Ohana (Cheng, Mailund, and R. Nielsen, 2017) with K=4 hidden ancestry clusters. We chose this value of K because, under this scheme, three of the components correspond to the three major ancestral populations that have been previously shown to have resulted - via multiple migration and admixture events - into the present-day European gene pool: the original Mesolithic hunter-gatherers (HG), the Neolithic farmers that migrated from the Near East (NEOL), and the Yamnaya steppe peoples that entered Europe during the Bronze Age (YAM) (Lazaridis, Patterson, et al., 2014; Allentoft et al., 2015; Haak et al., 2015) (Figure 1). The fourth is an ancestry component that remains largely confined to Northern Africa throughout most of the Holocene (NAF). In the following, we largely focus on the first three components and we note that the specification of a larger number of ancestry components could provide further details into more subtle patterns of migration and population expansion, but may also be confounded by bottlenecks and ghost admixture events (Daniel J Lawson, Van Dorp, and Falush, 2018), so we do not pursue finer ancestry estimation here.

**Figure 1:**
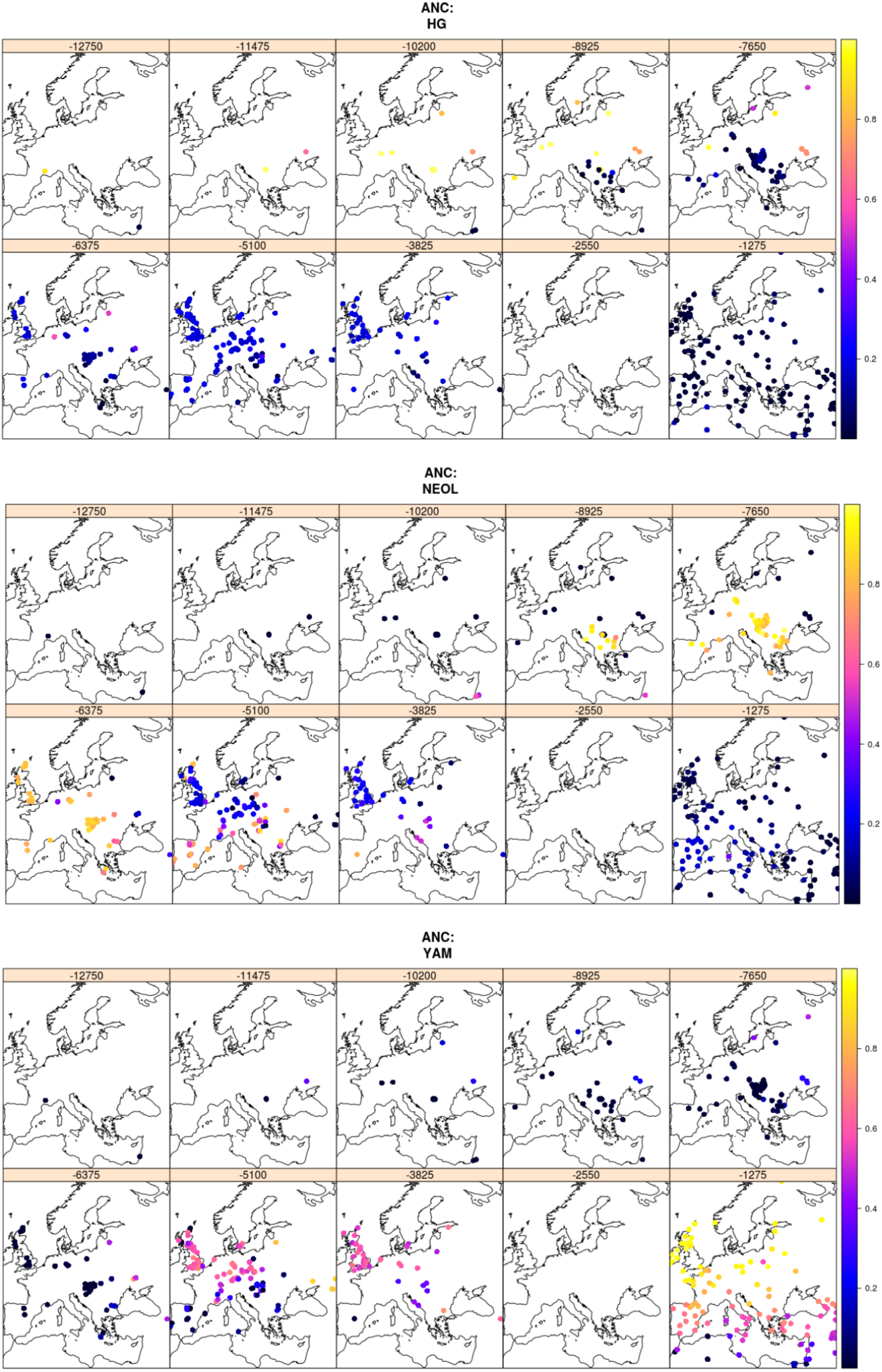
Spatiotemporal maps of ancestry proportions for ancient and present-day genomes in this study. The year in each panel’s title is the year of the most ancient sample in each panel, so not all ancient samples in each panel are strictly contemporaneous with each other. ANC:HG = hunter-gatherer ancestry. ANC:NEOL = Neolithic farmer ancestry. ANC:YAM = Yamnaya steppe ancestry.

As we show below, the YAM and NEOL ancestries closely parallel the Yamnaya and Neolithic farmer cultural horizons. However, ancestry and culture are distinct concepts that do not always overlap in time and space, so we choose to use the acronym nomenclature when referring to ancestries, and the full name when referring to cultures, unless otherwise specified. Furthermore, there were various, quite differentiated hunter-gatherer populations (Eastern, Western and Scandinavian hunter-gatherers) that migrated into Western Eurasia before the Holocene (Skoglund, Malmström, Omrak, et al., 2014; Skoglund, Malmström, Raghavan, et al., 2012; Sánchez-Quinto et al., 2012; Lazaridis, Nadel, et al., 2016; Fu et al., 2016). The HG ancestry roughly corresponds to the ancestry referred to as “Western hunter-gatherer" in these publications. However, the data for each of these populations is scarcer than for Bronze Age and Neolithic individuals, and, in this work, we chose only to focus on Holocene ancestry movements.

We first sought to compare the spread of dispersal of NEOL and YAM ancestries over time, using the calibrated C14 dates of each genome. Following Pinhasi, Fort, and Ammerman (2005) and Silva and Steele (2014), we regressed time against distance from the presumed origin of the spread of each of these ancestries, using the ranged major axis (RMA) method. This allows us to obtain an estimate for the migration front speed. We first used samples that had at least 50% of the corresponding ancestry we were studying, and then used a higher ancestry cutoff in which we only looked at samples with at least 75% of the corresponding ancestry. For the 50% ancestry approach, we find that the speed of the YAM migration (4.2 km/year; CI: 3.5-5.2) was at least twice as fast as the NEOL migration (1.8 km/year; CI: 1.6-2.1), assuming an origin of the Yamnaya migration at the center of its historical range (Figure 2.A). The higher ancestry cutoff approach yields the same estimate for the NEOL migration, but an even faster estimate for the YAM migration (9.4 km/year; CI: 6.1-20). Given that the original Yamnaya range was quite large (Anthony, 2010), we also aimed to see how our estimates varied as we varied the point of origin within the Yamnaya range. We obtained estimates of speed assuming a location of origin at the northern-most, eastern-most, western-most and southern-most parts of the Yamnaya range, which yielded similar estimates of speed (Table 1)). The correlation coefficients between time and distance from origin can also be used to estimate the point of origin (Pinhasi, Fort, and Ammerman, 2005; Hunt et al., 2018), assuming a range expansion for the YAM and NEOL ancestries. Indeed, when we varied the point of origin, we find that the most negative correlation coefficients correspond to Anatolia and the Middle East for the NEOL ancestry and to the Caspian steppe for the YAM ancestry (Figure 2.B).

**Table 1:**
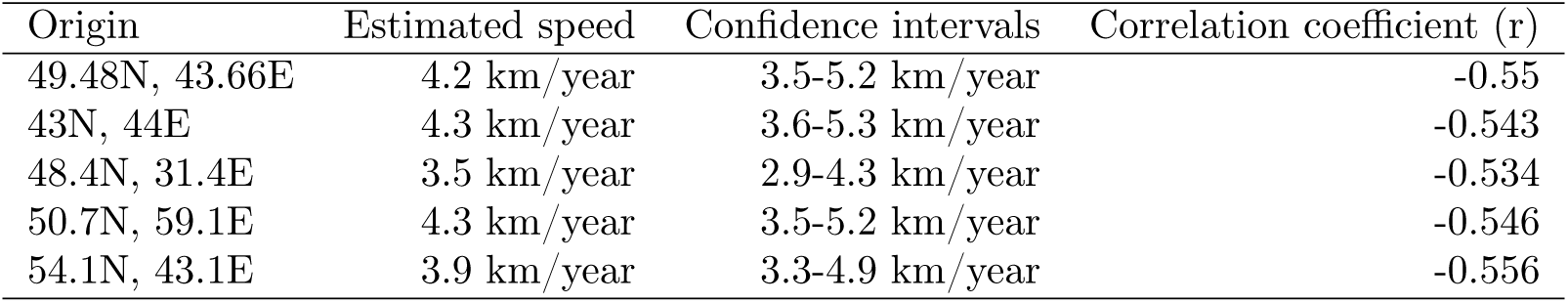
Estimated speed of the YAM ancestry expansion based on the genomes that have more than 50% of this ancestry, obtained by taking the negative inverse of the slope of an RMA regression of the point of origin against the time of sampling. We tried different points of origin at the center and extremes of the Yamnaya historical range. We also list the correlation coefficients obtained from the RMA analysis.

**Figure 2:**
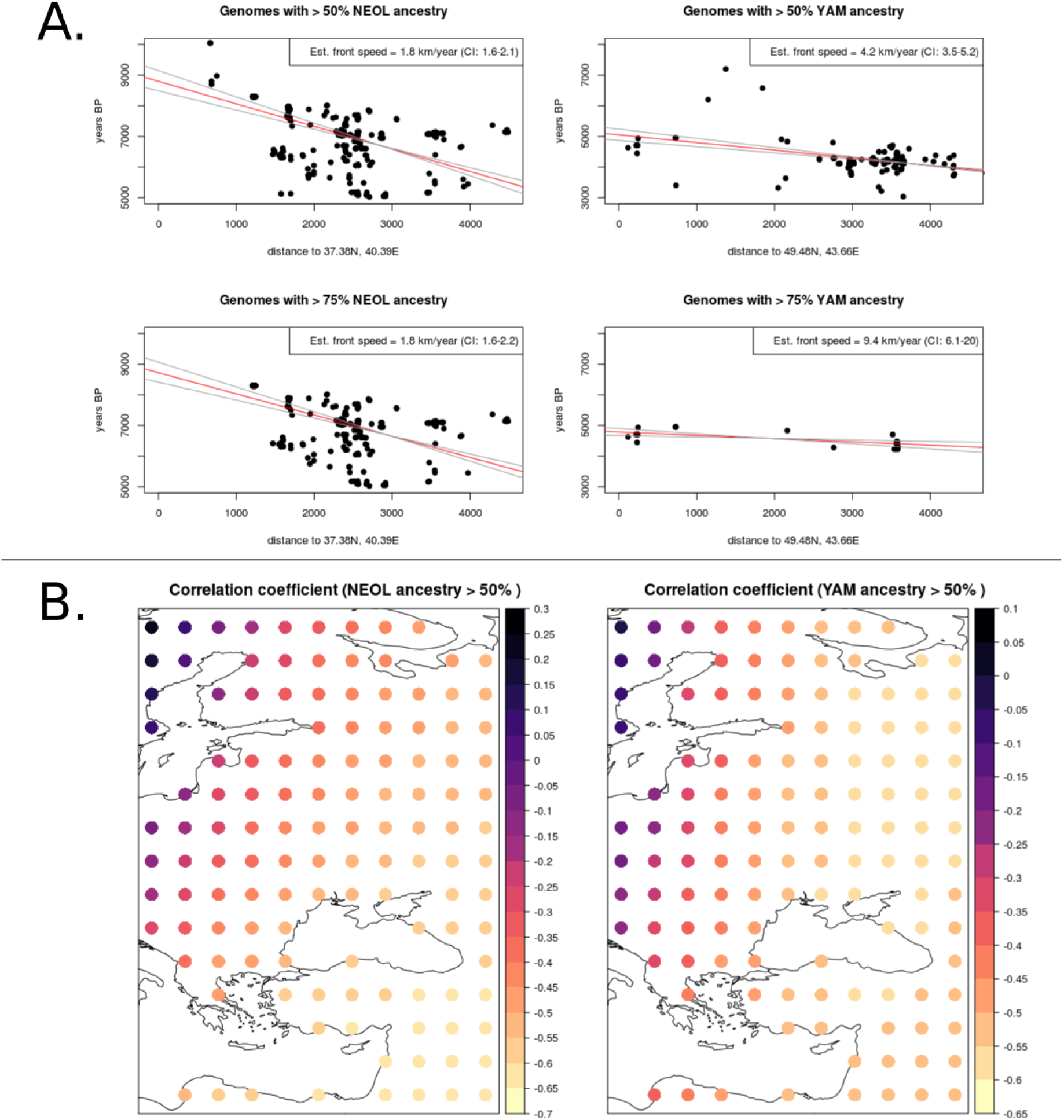
A. Front speed estimation for the Neolithic farmer (left column) and Yamnaya steppe peoples (right column) population movements. We used an RMA regression on time against distance from the hypothesized origin of the spread, to estimate average migration front speed. We used a lenient (*>* 50%) ancestry cutoff to define genomes as belong to a particular migration wave (top row) and a more conservative (*>* 75%) ancestry cutoff (bottom row). B. Point of origin estimation. We computed the correlation coefficient between time of sampling and distance from a hypothesized origin, which should be negative for a range expansion. Each dot in the map represents a different hypothesized origin.

To be able to compare ancestry across time and space with other variables, we aimed to project our ancestry values to particular times and locations for which we do not necessarily have sampled genomes (Figure 3). To do so, we computed a spatiotemporal variogram and fitted it to a metric covariance function (E. J. Pebesma, 2004; E. Pebesma and Heuvelink, 2016) (Figures S1,S2,S3,S4). Then, we performed spatiotemporal kriging of the inferred latent ancestry values on a dense grid of spatial points across Europe, over a 10,800-year span, with intervals of 400 years (Figures 4, 5, S6, S5, Supplementary Animations 1-3). In practice, however, given the sparseness of the data in the distant past, we restrict our discussion to patterns seen after 8,000 years BP.

**Figure 3:**
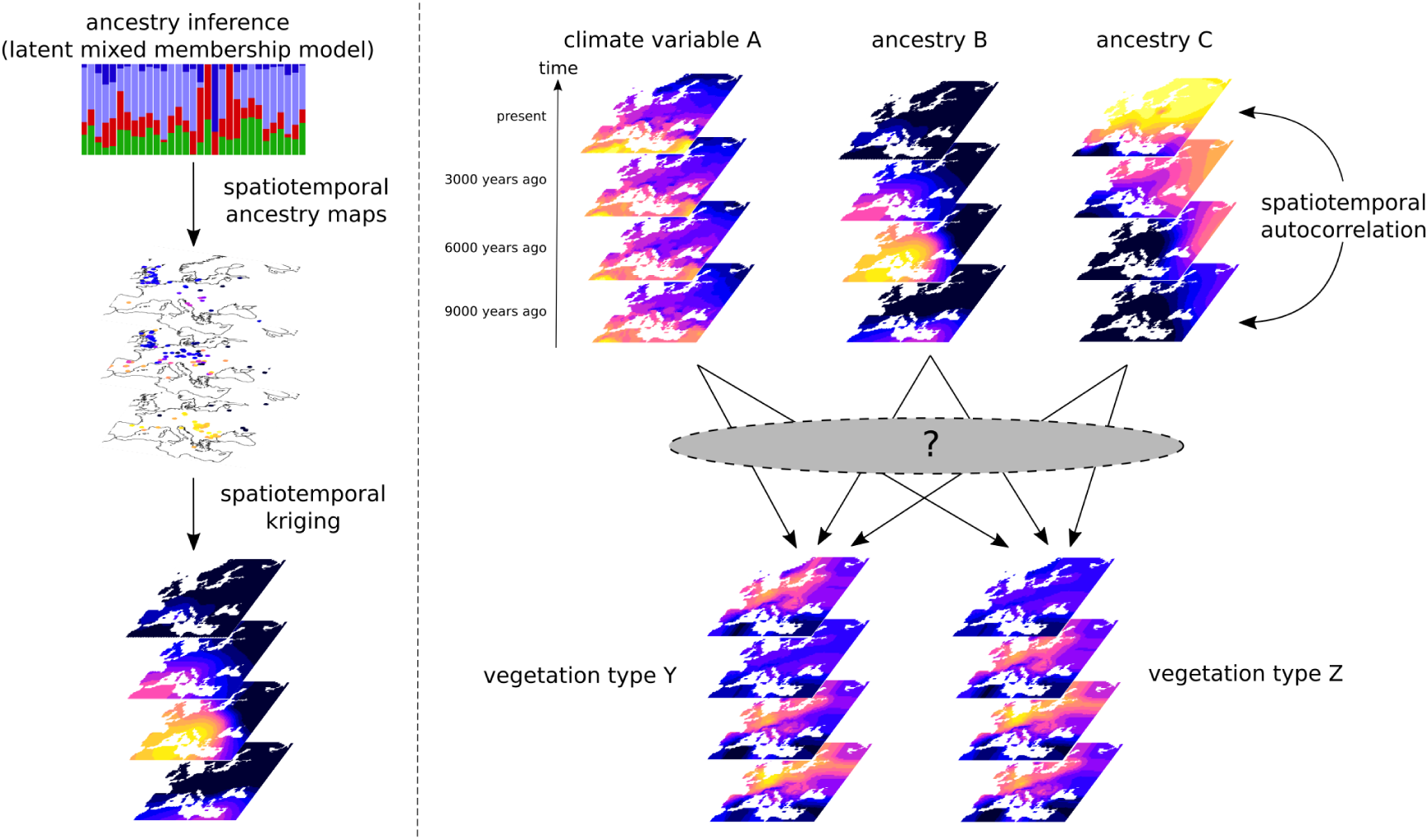
Schematic of methodology for spatio-temporal kriging and vegetation modelling. Left panel: We first fitted a latent mixed membership model to the ancient and present-day genomes. The ancestry proportions are then assigned the temporal and spatial meta-data of their respective genomes, which allows us to perform spatio-temporal kriging to any location and time in the European Holocene. Right panel: We used a spatiotemporally-aware model to understand how patterns of human migration and climate relate to patterns of vegetation type changes during the European Holocene, while accounting for spatiotemporal autocorrelation. We used a bootstrapping method to account for biases due to uneven sampling of ancient genomes. Brighter colors represent higher values of each depicted variable.

We also downloaded land cover class (LCC) maps (R. M. Fyfe, Woodbridge, and N. Roberts, 2015) and paleo-climatic variable maps (Brown et al., 2018) spanning the Holocene, and projected them on the same spatiotemporal grid that we used for our kriged ancestry values (see Methods). We computed correlations between each of the spatiotemporally projected ancestry proportions and vegetation types, and between the climate variables and vegetation types. This was done in three different ways. In the first, we simply obtained the correlation of the raw values of any two variables (Figure S7). In the second, we obtained the correlation of the differences in these variables at each time slice (Figure S9). In the third, we obtained the correlations of the variable anomalies, defined as the value of each ancient variable after subtracting the present-day value from the same location (Figure S8). We note, however, that this approach does not account for the autocorrelation in time and space that necessarily exists for all the compared variables, not only because of real autocorrelation in the processes under study, but also because of enforced autocorrelation from the smoothing techniques that generated the maps.

**Figure 4:**
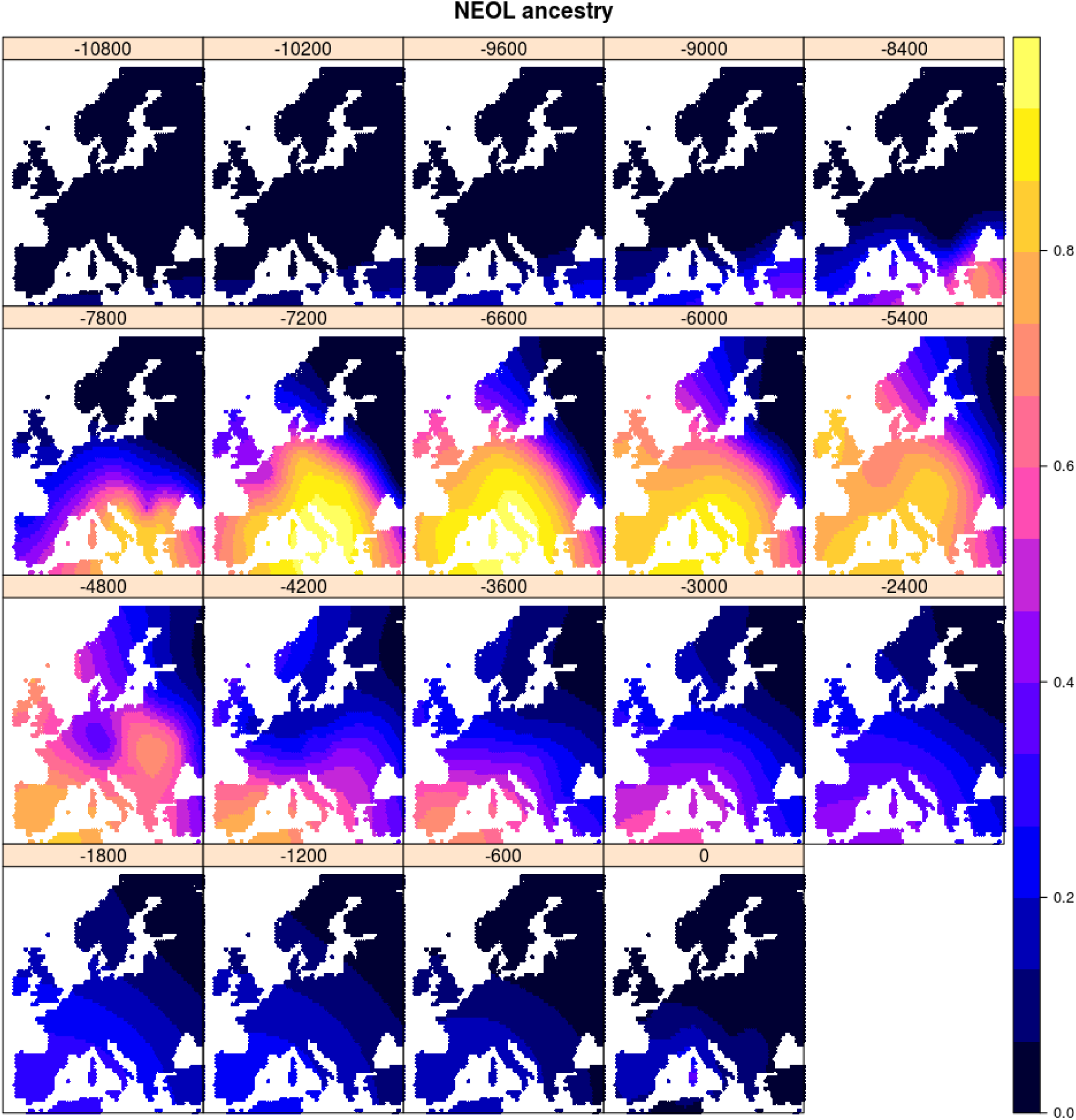
Spatiotemporal kriging of NEOL ancestry during the Holocene, using 5000 spatial grid points. The colors represent the predicted ancestry proportion at each point in the grid.

**Figure 5:**
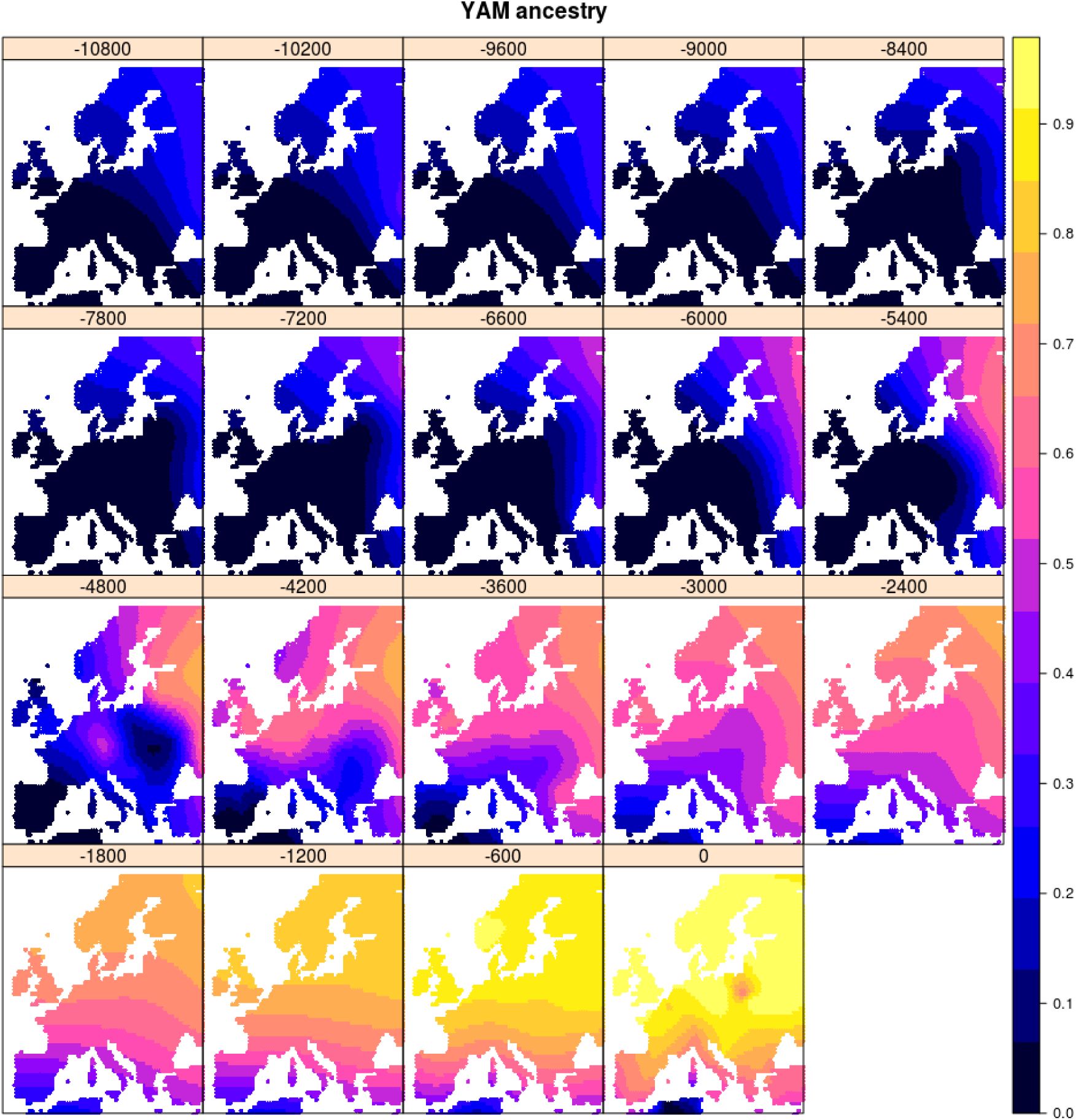
Spatiotemporal kriging of YAM steppe ancestry during the Holocene, using 5000 spatial grid points. The colors represent the predicted ancestry proportion at each point in the grid.

The raw correlations largely reflect spatially static patterns of co-occurrence (Figure S7, Table 2). For example, YAM ancestry is largely prevalent in northeast Europe throughout much of the Holocene, and this coincides with the prevalence of needle-leaf forests, which is why there is strong positive correlation between these variables. Conversely, NEOL ancestry is largely prevalent in southern Europe during this period, which is why there is a negative correlation with needle-leaf forest.

**Table 2:** Correlations between kriged ancestries and paleo-vegetation variables. Raw = correlation of raw variables corresponding to each spatio-temporal grid point. Differences = correlation of differences in raw variables at the same spatial grid point, between consecutive time slices. Anomalies = correlation in anomalies in variables, computed by subtracting the raw variable from its present-day value at the same spatial grid point.

| Correlation | Raw | Differences | Anomalies |
| --- | --- | --- | --- |
| NEOL vs. NEEDLE-LEAF FOREST | -0.473 | 0.0182 | 0.000753 |
| NEOL vs. BROAD-LEAF FOREST | -0.153 | 0.0392 | -0.065 |
| NEOL vs. HEATH / SCRUBLAND | 0.362 | 0.0135 | 0.152 |
| NEOL vs. PASTURE / NATURAL GRASSLAND | 0.35 | -0.0859 | -0.0777 |
| NEOL vs. ARABLE / DISTURBED LAND | 0.261 | -0.0978 | -0.0472 |
| HG vs. NEEDLE-LEAF FOREST | 0.176 | 0.0566 | -0.000407 |
| HG vs. BROAD-LEAF FOREST | 0.266 | -0.0356 | 0.212 |
| HG vs. HEATH / SCRUBLAND | -0.175 | -0.0762 | -0.176 |
| HG vs. PASTURE / NATURAL GRASSLAND | -0.172 | 0.0723 | -0.0527 |
| HG vs. ARABLE / DISTURBED LAND | -0.28 | 0.0688 | -0.113 |
| YAM vs. NEEDLE-LEAF FOREST | 0.439 | -0.0148 | -0.109 |
| YAM vs. BROAD-LEAF FOREST | 0.0806 | -0.098 | -0.503 |
| YAM vs. HEATH / SCRUBLAND | -0.279 | 0.0208 | 0.221 |
| YAM vs. PASTURE / NATURAL GRASSLAND | -0.352 | 0.12 | 0.467 |
| YAM vs. ARABLE / DISTURBED LAND | -0.22 | 0.0558 | 0.343 |

In contrast, the correlations in differences and in anomalies reflect spatially dynamic patterns of co-occurrence (Figures S9 and S8, Table 2). Here, temporal increases in one variable that coincide with temporal increases in a second variable at the same location will result in positive correlation, The same will result if there are co-occurring decreases. If, however, a variable decreases while another increases at the same location, this will result in negative correlation. For example, we see that YAM ancestry anomalies are positively correlated with pasture land anomalies, but negatively correlated with broad-leaf forest anomalies.

Focusing on particular locations of our map, we can better understand the reason for some of these correlations (Figure 6). For example, in central France, we observe that increases in YAM ancestry coincided with decreases in broad leaf forest cover, and, later on, with increases in pasture land-cover. This process was not spatially homogeneous, however. In southeastern and southwestern Europe, forest cover remained stable at low levels, even as YAM ancestry was increasing, perhaps due to the development of tree cropping within the agropastoral system in the Mediterranean (C. N. Roberts et al., 2019). Considerable increases in arable land-cover occur fairly late throughout the continent, and much later than the incursion of NEOL ancestry during the Neolithic. Interestingly, we observe a decrease in NEOL ancestry that continues even after the incursion of YAM ancestry into Europe, though our resolution for quantifying changes in this ancestry in the last 2,000 years is limited by the scarcity of ancient genomes from the very recent past.

**Figure 6:**
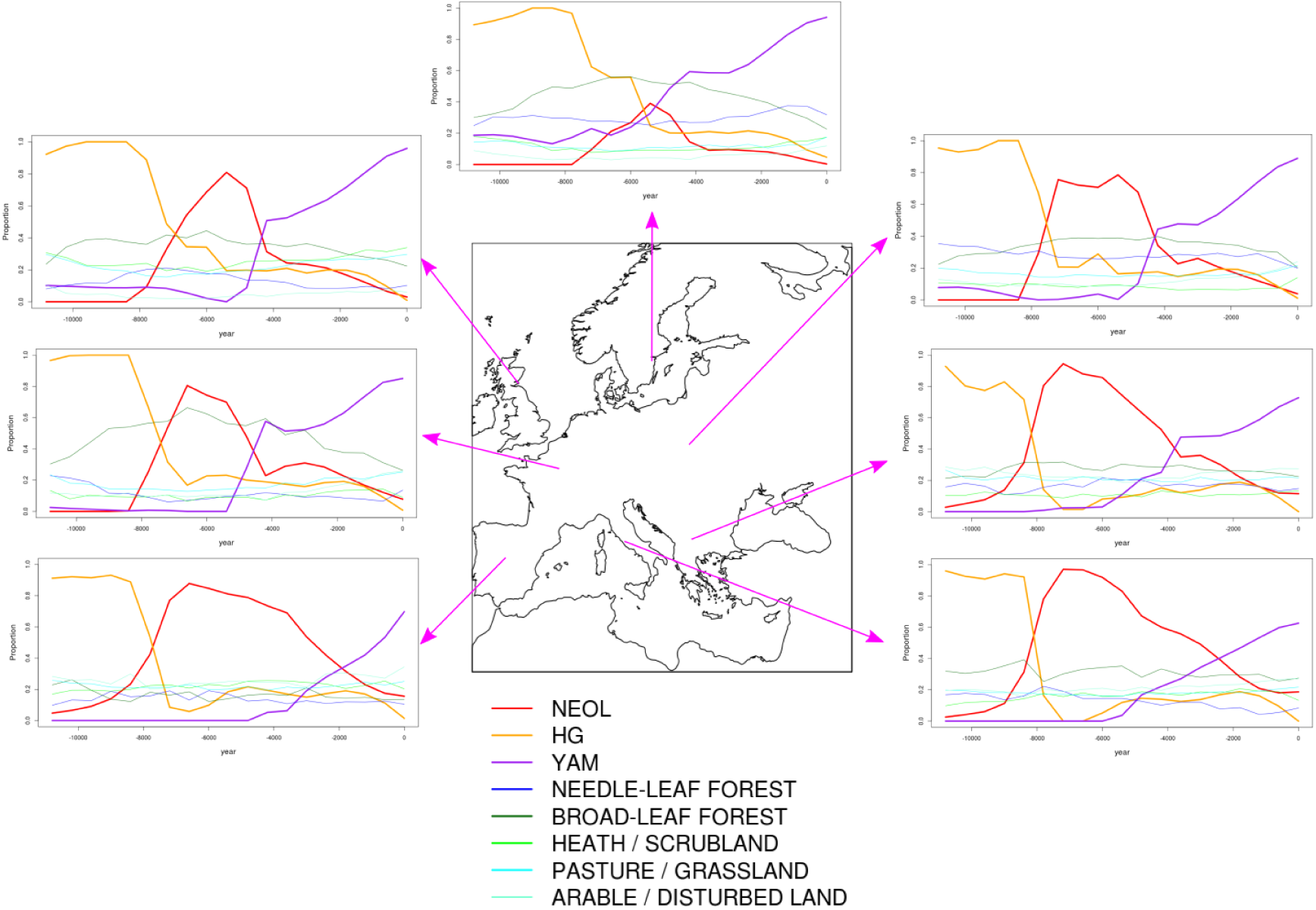
Timelines of kriged ancestry and vegetation type proportions at different points in Europe. NEOL = Neolithic farmer ancestry. HG = Hunter-gatherer ancestry. YAM = Yamnaya steppe ancestry.

While these correlation patterns are interesting, they do not account for the fact that we are projecting the data to lie in a particular set of grid points and time slices, which generate complex auto-correlations in time and space, potentially affecting the correlations we observe between variables. To address this, we used a spatiotemporally explicit hierarchical Bayesian model to better understand the relationships between changes in climate, ancestry and paleo-vegetation, while accounting for their autocorrelation in time and space (Figure 3). We used two models, implemented in the R package spTimer (Bakar, Sahu, et al., 2015). One is a Gaussian process (GP) model that incorporates a spatiotemporal nugget that is independent of time and has a distribution that depends on a spatial correlation matrix (Figure 7, Table 3). The other is an extension of this method that incorporates a temporal autoregressive component (AR; Figure 7, Table 4). We set the kriged ancestry and climate variables to be the explanatory variables, while each of the paleo-vegetation variables was set as a response variable. We fitted five separate models for each paleo-vegetation variable.

**Table 3:** Coefficients of the spatiotemporal Gaussian process model, using the PaleoClim (Brown et al., 2018) simulation-based paleo-climate variables as covariates, with 95% Bayesian credible intervals.

| Coefficient | Posterior median | 2.5% lower CI | 97.5% upper CI |
| --- | --- | --- | --- |
| NEOL → NEEDLE-LEAF FOREST | -0.0567 | -0.149 | 0.035 |
| HG → NEEDLE-LEAF FOREST | -0.196 | -0.329 | -0.0654 |
| YAM → NEEDLE-LEAF FOREST | -0.114 | -0.183 | -0.0434 |
| NEOL → BROAD-LEAF FOREST | -0.0396 | -0.113 | 0.033 |
| HG → BROAD-LEAF FOREST | 0.245 | 0.138 | 0.344 |
| YAM → BROAD-LEAF FOREST | -0.191 | -0.244 | -0.139 |
| NEOL → HEATH / SCRUBLAND | 0.0545 | -0.0584 | 0.167 |
| HG → HEATH / SCRUBLAND | 0.0426 | -0.114 | 0.2 |
| YAM → HEATH / SCRUBLAND | 0.0534 | -0.0262 | 0.132 |
| NEOL → PASTURE / NATURAL GRASSLAND | 0.0481 | -0.0026 | 0.0992 |
| HG → PASTURE / NATURAL GRASSLAND | 0.0196 | -0.0527 | 0.0947 |
| YAM → PASTURE / NATURAL GRASSLAND | 0.318 | 0.28 | 0.356 |
| NEOL → ARABLE / DISTURBED LAND | -0.0603 | -0.152 | 0.0275 |
| HG → ARABLE / DISTURBED LAND | -0.154 | -0.282 | -0.0285 |
| YAM → ARABLE / DISTURBED LAND | 0.0455 | -0.016 | 0.112 |

**Table 4:** Coefficients of the spatiotemporal autoregressive model, using the PaleoClim (Brown et al., 2018) simulation-based paleo-climate variables as covariates, with 95% Bayesian credible intervals.

| Coefficient | Posterior median | 2.5% lower CI | 97.5% upper CI |
| --- | --- | --- | --- |
| NEOL → NEEDLE-LEAF FOREST | -0.0114 | -0.0712 | 0.0492 |
| HG → NEEDLE-LEAF FOREST | -0.00947 | -0.0938 | 0.0801 |
| YAM → NEEDLE-LEAF FOREST | -0.00891 | -0.0533 | 0.0343 |
| NEOL → BROAD-LEAF FOREST | 0.324 | -0.032 | 0.105 |
| HG → BROAD-LEAF FOREST | 0.225 | 0.121 | 0.331 |
| YAM → BROAD-LEAF FOREST | -0.00494 | -0.0462 | 0.0361 |
| NEOL → HEATH / SCRUBLAND | -0.0289 | -0.0929 | 0.0361 |
| HG → HEATH / SCRUBLAND | -0.105 | -0.199 | -0.0107 |
| YAM → HEATH / SCRUBLAND | -0.00871 | -0.0556 | 0.0374 |
| NEOL → PASTURE / NATURAL GRASSLAND | -0.0271 | -0.0607 | 0.00651 |
| HG → PASTURE / NATURAL GRASSLAND | -0.153 | -0.207 | -0.0994 |
| YAM → PASTURE / NATURAL GRASSLAND | 0.0436 | 0.0219 | 0.0657 |
| NEOL → ARABLE / DISTURBED LAND | -0.0392 | -0.0827 | 0.0043 |
| HG → ARABLE / DISTURBED LAND | -0.0831 | -0.146 | -0.0208 |
| YAM → ARABLE / DISTURBED LAND | -0.0111 | -0.0415 | 0.0212 |

**Figure 7:**
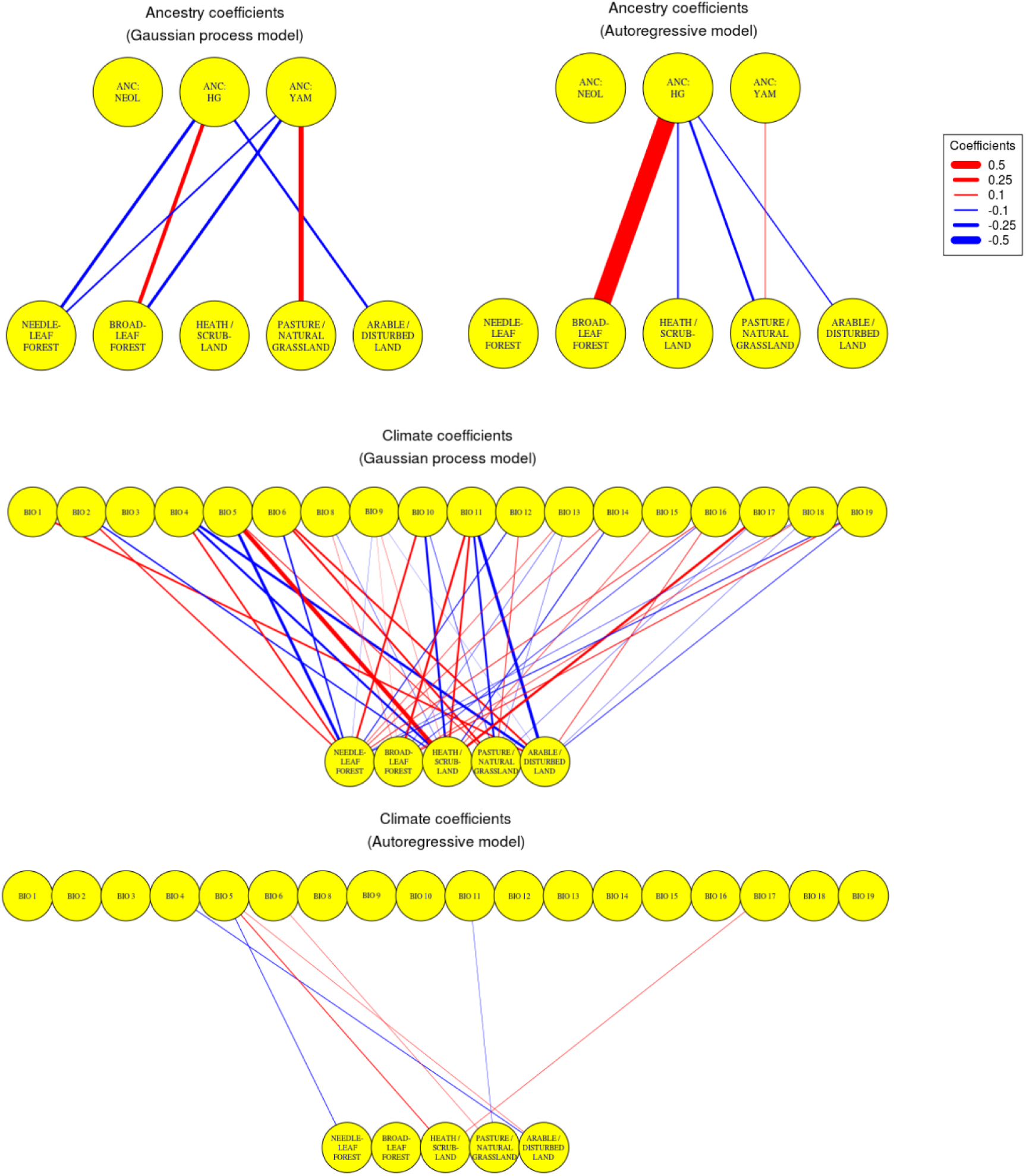
Posterior mean coefficients of spatiotemporal model for paleo-vegetation anomalies, using kriged ancestry anomalies and anomalies from simulation-based paleo-climate reconstructions as explanatory variables. Upper left and middle panels: posterior coefficients from Gaussian process model. Upper right and bottom panels: coefficients from autoregressive model. Coefficients whose corresponding posterior distribution has a 95% central probability mass interval that spans the value of 0 are not depicted. ANC:NEOL = neolithic farmer ancestry. ANC:HG = hunter-gatherer ancestry. ANC:YAM = Yamnaya steppe ancestry. The climate variables follow the WorldClim nomenclature. BIO1 = Annual Mean Temperature. BIO2 = Mean Diurnal Range (Mean of monthly (max temp - min temp)). BIO3 = Isothermality (BIO2/BIO7) (* 100). BIO4 = Temperature Seasonality (standard deviation *100). BIO5 = Max Temperature of Warmest Month. BIO6 = Min Temperature of Coldest Month. BIO8 = Mean Temperature of Wettest Quarter. BIO9 = Mean Temperature of Driest Quarter. BIO10 = Mean Temperature of Warmest Quarter. BIO11 = Mean Temperature of Coldest Quarter. BIO12 = Annual Precipitation. BIO13 = Precipitation of Wettest Month. BIO14 = Precipitation of Driest Month. BIO15 = Precipitation Seasonality (Coefficient of Variation). BIO16 = Precipitation of Wettest Quarter. BIO17 = Precipitation of Driest Quarter. BIO18 = Precipitation of Warmest Quarter. BIO19 = Precipitation of Coldest Quarter.

In comparison to the GP model, the AR model results in a more sparse set of posterior coefficients whose confidence intervals do not overlap with 0 (Figure 7, Table 4). An evaluation of the root mean squared error of the predictions (see Methods) suggests that the GP model with a uniform prior distribution for the spatial decay parameter of the Matérn correlation function generally has the best predictive accuracy (Figure S11). We also compared the predictive accuracy of models incorporating ancestry only, climate only, both sets of variables or none of them. Our results suggest that, for almost all the paleo-vegetation variables, there is no observable difference in predictive power in adding climate or ancestry, with the exception of pasture (LCC6), which shows the lowest predictive error when including both climate and ancestry under the GP model (Figure S11).

Because we are using kriged ancestry as an explanatory variable and our ancient genomes are unevenly sampled across space and time, we were mindful that the Bayesian credible intervals (BCI) obtained from the hierarchical model would not accurately reflect uncertainty in particular regions of space-time. For that reason, we performed nonparametric bootstrapping of the parameter estimates. We randomly sampled ancient and present-day genomes with replacement from among the list of all genomes until we had as many genomes as were in the original dataset, then obtained their ancestry assignments, kriged them on the spatiotemporal grid and inputted them into the Bayesian hierarchical model. We did this 100 times to obtain 100 pseudo-samples, which allowed us to obtain 95% bootstrap-based confidence intervals (BBCI) around the mean posterior estimates (Figure S10, Table 5). Below, we discuss results that are supported by one or both models, and that are also supported by the bootstrapping approach.

**Table 5:**
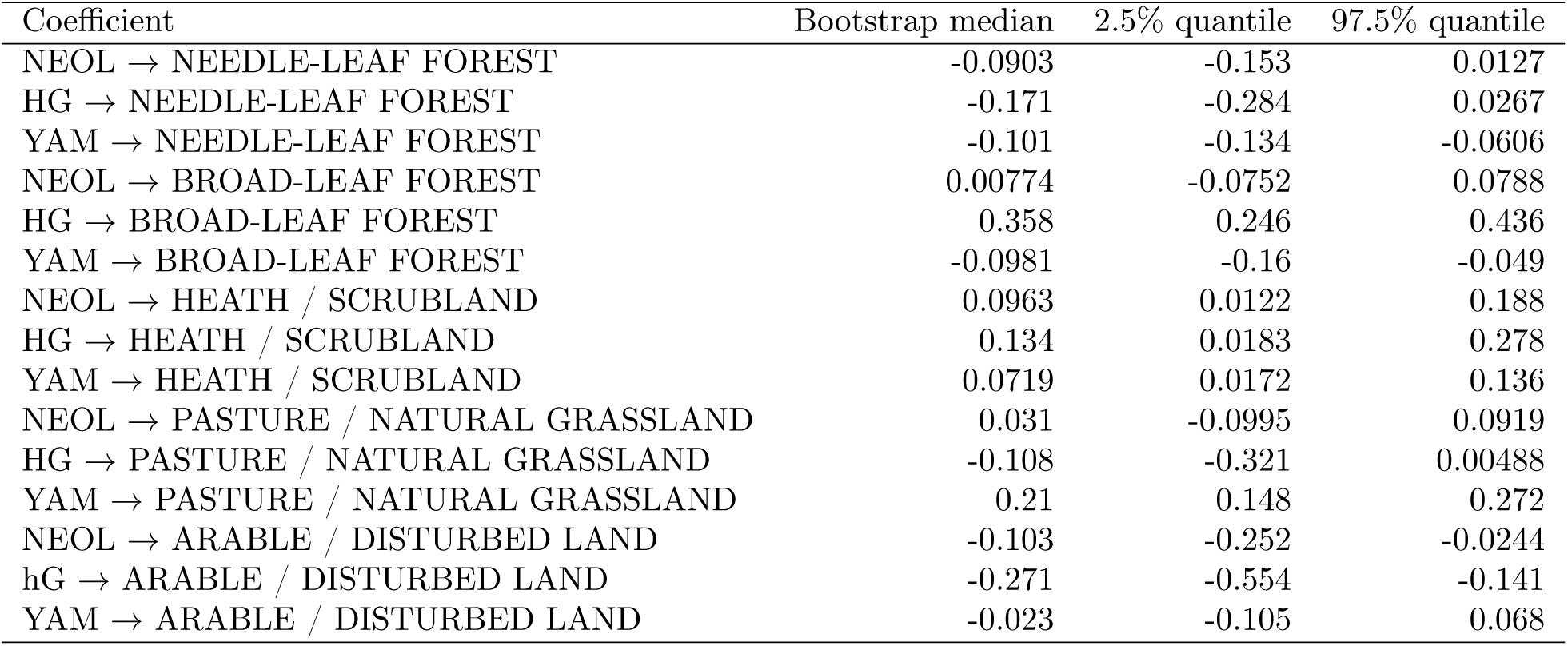
Coefficients of the spatiotemporal autoregressive model, using the PaleoClim (Brown et al., 2018) simulation-based paleo-climate variables as covariates, with 95% bootstrap-based confidence intervals.

Regardless of the model used, we find that HG ancestry contributes positively to broad-leaf forest anomalies, but negatively to arable land anomalies. YAM ancestry, in turn, contributes positively to pasture (Figures 7, S10). In the fitted AR model and the bootstrapping approach, we also see a negative contribution of HG ancestry to pasture and scrubland. In the fitted GP model and the bootstrapping approach, we observe a negative contribution of YAM ancestry to forest vegetation, which is strongest for broad-leaf forest. We see weak or non-existent contributions of NEOL ancestry to any vegetation type, suggesting that the population movement associated with NEOL ancestry did not have an immediate effect on vegetation composition, or that the effect was not large enough to be detected. We cannot discard the possibility that we may lack the power to detect some of these changes at our current scale of resolution.

We additionally observe non-zero contributions of different climate variables to the different vegetation anomalies, which become sparser in the AR model (Figure 7). For example, in both the AR and GP models, increases in temperature are related to increases in non-forest vegetation types (scrubland, pasture and arable land). In addition, temperature seasonality may be interpreted as contributing negatively to the proportion of arable land, while precipitation during the driest quarter may be interpreted as contributing positively to heath / scrubland, under the fitted model. Finally, we also built “first arrival” maps (Pinhasi, Fort, and Ammerman, 2005; Fort, 2015; Vander Linden and Silva, 2018) for both the NEOL ancestry and the YAM ancestry, given that these ancestry changes can be broadly interpreted as incursions of foreign populations into the European continent during the Neolithic and Bronze Age (Allentoft et al., 2015; Haak et al., 2015). Here, we record the approximate period of first appearance of individuals with more than some particular ancestry proportion cutoff (Figure 8). Constructing this map for NEOL ancestry allows us to observe that this ancestry spread closely parallels the inferred cultural spread of farming, which has been inferred from archaeological sites (Fort, 2018; Fort, 2015; Pinhasi, Fort, and Ammerman, 2005; Silva and Steele, 2014). When performing the same type of reconstruction for the YAM ancestry, we observe that this spread occurs first via north and central Europe, and only much later begins to spread into southern Europe (Figure 8), again reflecting reconstructions from archaeological records for the spread of the Yamnaya, Corded Ware and Bell Beaker cultures.

**Figure 8:**
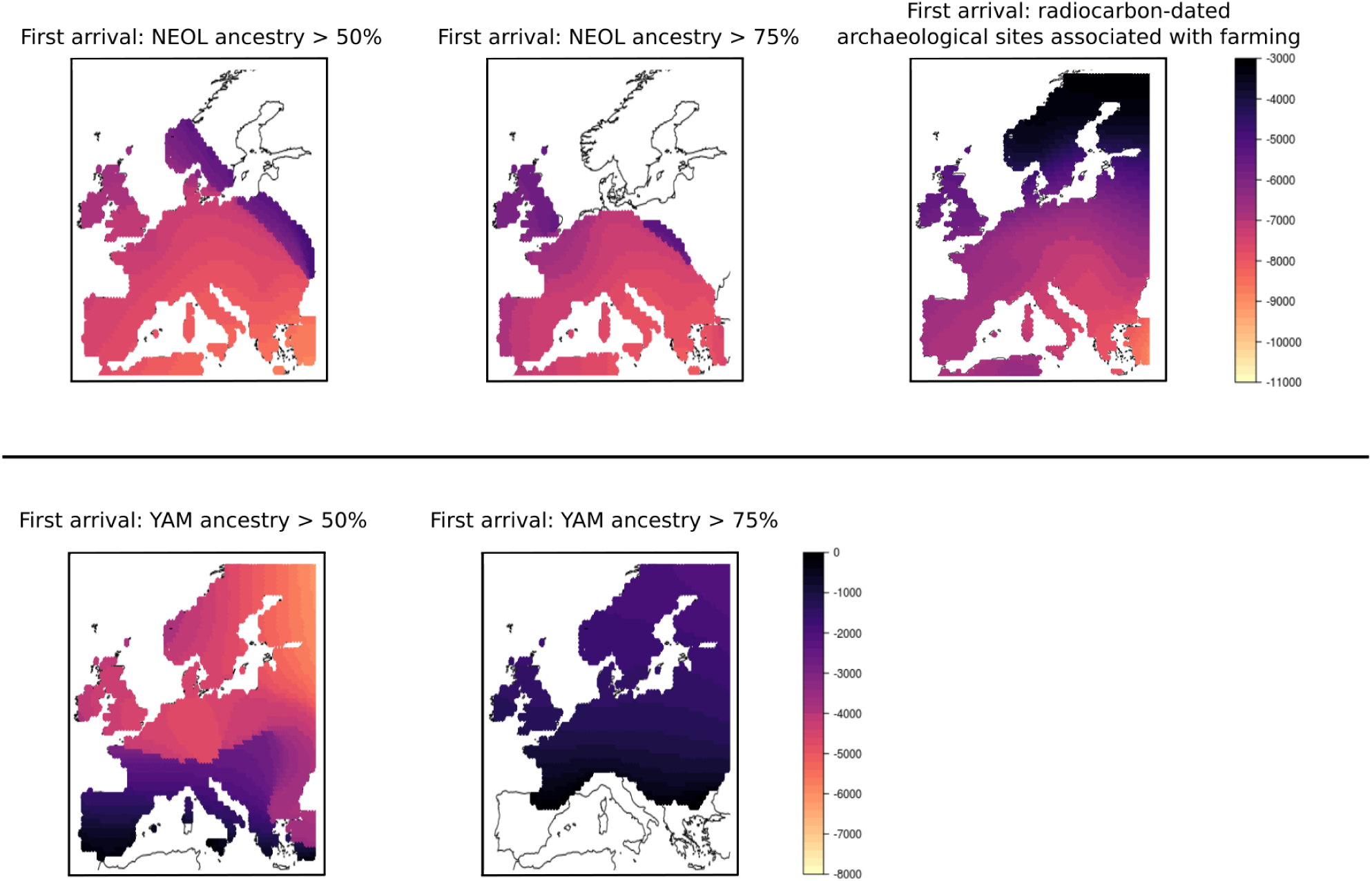
Comparison of inferred spread of farming from archaeological sites and spread of NEOL (upper panel) and YAM ancestries (lower panels). The left panels define first arrival as the first time slice in which a grid point has more than 50% of the ancestry depicted. The center panels are the result of using a more strict cutoff: 75% ancestry. The top right panel is a spatially kriged map of first arrivals of farming practices, based on radiocarbon-dated archaeological sites. White regions are either regions without any genomes included in our data or regions that do not contain genomes passing the assigned cutoff throughout the duration of the Holocene.

## Discussion

An explicitly geostatistical approach allows us to easily visualize how movements of ancestry occurred during the Holocene in Europe. We can observe, for example, that the NEOL ancestry expansion followed a two-pronged shape, paralleling the expansion of farming practices estimated from radiocarbon-dated archaeological sites. In particular, we observe two wave fronts, one northward across central Europe, and one westward along the Mediterranean coast (Figure 8). In the cultural map, these two wave fronts correspond to the Linear Pottery culture (LBK) (Bickle and Whittle, 2013) and the Impressa / Cardial Pottery culture (Barnett, 2000; Binder et al., 2018; Manen et al., 2019). Given their close parallels in the ancestry map, this supports the view that these two cultural expansions may have been driven by migrations of people (Allentoft et al., 2015; Haak et al., 2015).

We estimate that the expansion of YAM ancestry occurred faster than the expansion of NEOL ancestry. The reasons for this could be numerous, including the use of horses for long-distance travel (Anthony, 2010). YAM ancestry predominates in individuals associated with the Yamnaya and Corded Ware cultures, and is presumed to have moved into Europe from the Eurasian Steppes (Kristiansen and Larsson, 2005). Another possibility could be the opening of the landscape previous to the arrival of the Yamnaya people, perhaps due to Neolithic agricultural, grazing and mining practices (Schauer et al., 2019), which may have facilitated later movements of people. We do not observe a strong decrease in forest vegetation in Northern and Central Europe until the Bronze Age, however. On the other hand, there is limited evidence that Corded Ware people were horse herders, and evidence from settlements in central Europe suggests they may have practiced mixed agriculture (Müller et al., 2009; Seregély and Müller, 2008; Jacomet, 2008).

This approach also allows us to model ancestry together with other variables for which there are rich spatio-temporal estimates, namely model-based paleo-climate and pollen-based vegetation landcover types, while accounting for the spatio-temporal autocorrelation among these variables (Figure 7). Our models incorporating both climate and ancestry generally fit HG ancestry as positively contributing to forest-type vegetation, while YAM ancestry as negatively contributing to this form of vegetation, and positively contributing to pasture and arable land.

We do not find that NEOL ancestry had much of a strong contribution to changes in vegetation, suggesting that this expansion did not have an immediate effect on the European landscape, or that this impact was too weak or localized for us to clearly detect an effect in our model. Earlier studies have shown that Neolithic communities did in fact alter their local environments (Mercuri et al 2019 Holocene) and had a local impact on vegetation to a certain extent, at least in northwestern Europe (Woodbridge, R. M. Fyfe, et al., 2014; Schauer et al., 2019; Lechterbeck et al., 2014; Woodbridge, R. M. Fyfe, et al., 2014). Figure 6 shows a very minor decline in broad-leaf forest that coincided with the increase in NEOL ancestry, in particular areas such as northern and northwestern Europe. However, a much more pronounced reduction in broad-leaf forest occurs later on throughout western and northwestern Europe, and coincides with the increase in YAM ancestry. In relation to this point, it is important to note that cultivated tree types (olive, chestnut and walnut) - which are pervasive in the Mediterranean - also fall into the category of broad-leaf forest. Thus, our capacity to infer causes for changes in forest types in regions with this type of cultivar (e.g. the Mediterranean, Woodbridge, N. Roberts, and R. Fyfe (2018)) is limited.

The decrease in broad-leaf forest (starting 6,000 years BP) was followed by a minor increase in pasture and disturbed land in some parts of the continent. These vegetation types are naturally present in the Mediterranean and the Black Sea region throughout the earlier part of the Holocene and remain fairly stable until the present (R. M. Fyfe, Woodbridge, and N. Roberts, 2015). In contrast, in western Europe, these vegetation types only reach intermediate levels during the Bronze Age - as YAM ancestry begins to increase - and they continue to increase after the end of this period. Furthermore, increases in YAM ancestry in southern and eastern Europe do not coincide with increases in pasture and disturbed land.

Pasture is the only land cover type whose predictive power considerably increases as a result of adding ancestry and climate variables into our model. This might be because we currently lack the spatiotemporal resolution to provide much predictive power with the addition of climate or ancestry variables, or because these variables may not be strongly predictive of the other land cover classes. Other unaccounted factors may have had a stronger effect on the vegetated landscape. An obvious candidate is the dramatic increase in population density that occurred over the last 3,000 years (Bevan et al., 2017), which likely led to strong changes in land use practices, consequently disturbing vegetation throughout the continent. Earlier population rises and collapses during the Neolithic and Bronze Age could have also influenced the vegetated landscape in significant ways, although on a smaller scale (Shennan et al., 2013; Lechterbeck et al., 2014; Woodbridge, R. M. Fyfe, et al., 2014). Thus, a future study could aim to incorporate estimates of human population density or other measures of human activity into explanatory models for changes in vegetation, together with population movement. A recent approach using human land use estimates, for example, showed that, on a continental scale, climate changes were the main driver of changes in Holocene vegetation, but the influence of human land use increased in the later stages of the period (Marquer et al., 2017).

There are a number of caveats and assumptions in our modelling procedure that are important to stress. First, we are assuming that changes in ancient ancestries can be used as a proxy for long-distance movement of people. This may be the case for particular periods of time - especially when peoples of highly divergent ancestries first met each other - but this assumption loses validity as we move closer to the present, and the ancestry components tend to become more homogenized due to later migrations within the area of study (Margaryan et al., 2019). Tracing relatively high YAM ancestry in the present day is approximately equivalent to tracing people with high Northern European ancestry, who cannot be equated with ancient “steppe peoples”.

Second, we are relying on existing ancient DNA data, which has its own idiosyncrasies, due to environmental and historical biases in sampling. For example, North Africa and eastern Scandinavia are sparsely sampled in our dataset, so our ancestry estimates for those regions are much poorer than for the rest of the European continent. We attempted to prevent this type of biases from affecting our model inference via a bootstrapping approach, to assess how robust these were to accidents of sampling.

Additionally, we are relying on a particular choice of the number of hidden clusters or components (K) under a latent mixed-membership model. We chose this model and parameter setting to be able to discretize patterns of ancestry into three major population clusters (HG, NEOL and YAM), which have been documented via other, more involved, population genetic analyses (Haak et al., 2015; Allentoft et al., 2015; Lazaridis, Patterson, et al., 2014). We also chose a low number of clusters to have enough data points across extended periods of time, in order to accurately estimate the space-time decay in covariance between ancestries (e.g. Figures S1 and S2). These clusters are, however, an approximation of a very complex genealogical process. Indeed, the clusters cannot be seen as discrete, originally isolated populations, as there may be both isolation-by-distance and hierarchical population structure within each of these groups (Frantz et al., 2009; Janes et al., 2017; Battey, Ralph, and Kern, 2019). The clusters themselves are also the result of more complex admixture and migration events that occurred before the Holocene (Fu et al., 2016). Additionally, there is likely a large degree of differentiation in population structure over time and space, even when looking at the same admixture components (e.g. the NEOL component of a present-day Sardinian is differentiated relative to the NEOL component of a Bronze Age central European). These subtle patterns are hard to pick up by simple latent mixed-membership models (Daniel J Lawson, Van Dorp, and Falush, 2018), although there has recently been some progress in this regard (Joseph and Pe’er, 2018; Bradburd, Coop, and Ralph, 2018). Other types of population genetic frameworks are able to better detect some of these more subtle signals by, for example, modelling patterns of haplotype sharing (Hellenthal et al., 2014; Daniel John Lawson et al., 2012), the full site-frequency spectrum (Excoffier et al., 2013; Kamm et al., 2019) or an approximation to the full ancestral recombination graph (Kelleher et al., 2018; Speidel et al., 2019). Nevertheless, these also have their own limitations and assumptions. For all these reasons, we advice the reader to consider that the ancestry components used in this study are approximations of the true historical admixture process.

An improvement to our current approach could involve developing a hierarchical dynamical model for explicitly modelling spatiotemporal movements on the genetic data directly, without relying on ancestry assignments estimated from a non-spatiotemporally-aware model (Cressie and Wikle, 2015). This could also help with better dealing with boundary constraints that are not accounted for by the kriging methodology. For example, we currently have to correct kriged estimates that are lower than 0 or higher than 1. A generative model of spatio-temporal ancestry would not allow for these types of parameters in the first place, for example, by placing Bayesian priors on ancestry with 0 probability outside of the 0-1 range. This could also be solved by extending compositional interpolation techniques to a spatiotemporal setting (Walvoort and Gruijter, 2001).

We believe there is also a lot of potential for new geostatistical approaches that could be deigned to combine other types of datasets in an integrative approach for the study of the past, even at more local scales than we looked at here. This could encompass, for example, the combination of strontium and oxygen isotope analyses together with radiocarbon data and contextual archaeological information (Sjögren, Price, and Kristiansen, 2016; Mittnik et al., 2019), the joint analysis of genetic and linguistic changes over time (Kristiansen, Allentoft, et al., 2017) or the study of the interactions between population density and vegetation (Müller, 2015; Kolář et al., 2018).

In summary, the methodologies used here rely on several assumptions, and could benefit from a myriad of extensions. However, accounting jointly for space and time in the study of genetic and environmental variables is a necessary first step towards better understanding the impact of major migrations in the past. Without explicitly modelling space and time, researchers might miss local phenomena that may have unravelled in different ways across the area of study. This could lead us to ignore important historical processes that become occluded by taking an overly global perspective. We should not ignore the forest for the trees, but sometimes, the trees themselves might be hidden by the forest.

## Methods

All R code used to perform the analyses in this manuscript has been deposited in: https://github.com/FerRacimo/STAdmix

### Kriged ancestry maps over time and space

For our ancestry analyses, we used a combined dataset of 842 ancient and 955 present-day genomic sequences. The present-day genomes were typed at the Human Origins SNP array, while the ancient genomes were either typed at this array or whole-genome sequenced, followed by filtering for SNPs that are in this array (Mathieson, Lazaridis, et al., 2015; Lazaridis, Patterson, et al., 2014; Patterson et al., 2012; Allentoft et al., 2015). We restricted our analyses to modern human genomes obtained from human remains located within an area encompassing most of the European continent: north of 30^◦^N, south of 75^◦^N, east of 15^◦^W and west of 45^◦^E. We inferred latent ancestry components on these genomes using Ohana (Cheng, Mailund, and R. Nielsen, 2017).

We performed ordinary global spatio-temporal kriging using the R libraries gstat (E. J. Pebesma, 2004; E. Pebesma and Heuvelink, 2016) and spacetime (E. Pebesma et al., 2012), to obtain unbiased linear predictions of ancestry for unsampled locations and times. Suppose we have a set of noisy observations of a variable distributed unevenly across space and time. In our case, this will be the inferred proportion of a particular ancestry in each of our ancient genomes. Let ***s***_***i***_ be a vector representing the *i*th site (out of *n*) in our grid, which is composed of two values: its longitude and latitude. Following the notation by Cressie and Wikle (2015), suppose we have *T*_*i*_ different temporal samples of a measured variable at site ***s***_***i***_. A temporal sample obtained at the *j*th time (*t*_*ij*_) from this site will be denoted as *Z*(***s***_***i***_, *t*_*ij*_). Suppose these data are equal to the true spatiotemporal process plus some measurement error *ϵ*:

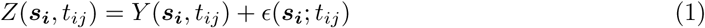

Let ***Z***^(*i*)^ be the vector containing all values that were measured at different time points in location *s*_*i*_. Also, let ***Z*** = (*Z*^(1)′^, …, *Z*^(*m*)′^)^′^ where *m* is the number of locations sampled. We can obtain a linear predictor, *Y* ^∗^(*s*_0_; *t*_0_) for a particular unsampled data point at time *t*_0_ and location *s*_0_:

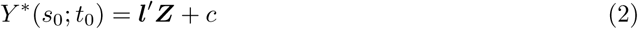

where ***l*** and *c* are parameters than can be optimized. In particular, for the case that the true process *Y* (;) has a constant unknown mean *µ*, one can show that the linear unbiased predictor that minimizes the mean squared prediction error - also called the ordinary kriging predictor - is equal to:

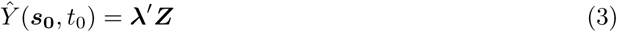

where 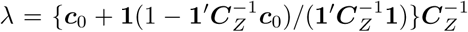, ***c***_0_ = *var*(***Z***) and ***C***_*Z*_ = *cov*(*Y* (***s***_0_, *t*_0_), ***Z***). The latter can be obtained by fitting a spatio-temporal covariance function for the true process ***Y*** to the empirical spatio-temporal variogram of the observed measurements ***Z*** (Figures S2, S1, S3, S4). In our case, the variogram was computed over a range of 3,000 years, with 60-year windows, and we used the “metric" variogram model to fit it (Gräler et al. 2015). For a more extensive explanation of spatiotemporal kriging, we refer the reader to Cressie and Wikle (2015).

As our predicted grid, we used a set of spatial points distributed evenly across Europe. We used two types of spatial grids: one containing a dense set (5000) of points and a sparser set, containing 200 points. We call this our "spatial grid". The dense version of the spatial grid was used for plotting spatiotemporal maps (e.g. Figure 5), while the sparse set was used to fit the Bayesian spatio-temporal model (e.g. Figure 7), for ease of computation. We observed that the ancestry-vegetation and climate-vegetation correlations computed under both schemes were almost identical, suggesting that the use of the sparser grid should not affect inference under the Bayesian model. Potential biases arising from particular grid points possessing few nearby ancient genomes are accounted for in the bootstrapping method described below. Unless otherwise stated, our "temporal" grid had a 10,800-year span, with intervals of 400 years until the present, for a total of 28 time slices. Thus, if our spatial grid had *a* spatial points, our "spatiotemporal grid" had 28*a* spatiotemporal points. We bounded the kriged ancestry values between 0 and 1, and so kriged values that were negative were set to 0, and those that were larger than 1 were set to 1.

In all analyses below, we did not include a kriged ancestry component that is largely restricted to north Africa (NAF). The reasons for this are two-fold: 1) this ancestry remains largely spatially static throughout the Holocene, at least with respect to the box we defined to bound our analyses; 2) given that all latent ancestries must add up to 1 in each individual genome, this ancestry is equal to 1 minus the sum of the other three ancestries, and is therefore not linearly independent from them.

### Paleo-vegetation maps

We downloaded inferred Holocene paleo-vegetation spatiotemporal maps from R. M. Fyfe, Woodbridge, and N. Roberts (2015). This paleo-vegetation reconstruction was built from 982 pollen records across Europe, using the pseudobiomization method (PBM) (R. Fyfe, N. Roberts, and Woodbridge, 2010). It has a 10,800-year span, with intervals of 200 years until the present. To ease computation, we sampled every two time windows, resulting in intervals of 400 years until the present, and for each paleo-vegetation time slice, we rasterized the map to have 6,540 points (down from 35,856). Then, for each time slice, we inferred the value of each point in our spatial grid by taking the median of the 5 nearest points in the rasterized map.

### Paleo-climate maps

We obtained a set of simulation-based Holocene paleo-climate reconstructions for Europe from PaleoClim (Brown et al., 2018), which includes surface temperature and precipitation estimates for the Early (11.7-8.326 kya), Middle (8.326-4.2 kya) and Late Holocene (4.2-0.3 kya), using snapshot-style climate model simulations. These simulations were accessed through PaleoView (Fordham et al., 2017) and come from the TRaCE21ka experiment (Liu, B. Otto-Bliesner, et al., 2009; Liu, Lu, et al., 2014), which used the Community Climate System Model v3 (CCSM3) (Bette L Otto-Bliesner et al., 2006; Collins et al., 2006; Yeager et al., 2006): a general circulation model involving atmosphere, ocean, sea ice and land. The PaleoClim authors refined the simulations from this model, incorporating small-scale topographic nuances of regional climatologies, thus creating high-resolution paleo-climate maps. We projected the three Holocene maps - together with the present-day WorldClim map - onto the previously delineated temporal grid, for each of the 19 climate variables that are present in the PaleoClim database. At each time slice, for each point in the our spatial grid, we inferred the value of each climate variable, by taking a weighted average of the values of the two closest bounding paleo-climate time points (past and future) at that spatial point, weighted by their respective temporal distance to our time slice. These allowed us to obtain a spatiotemporal grid of the climate variables at the same locations and times for which we had kriged ancestry and paleo-vegetation data. In the Bayesian hierarchical model, we excluded one of these variables (temperature annual range) because it is a linear combination of two of the other climate variables.

### Computation of correlations

We computed Pearson correlations between the kriged ancestry, climate and vegetation variables in three ways. First, we simply took the vector containing the values of one variable across all points in our spatiotemporal grid and computed its correlation with the values of another variable at all the same spatiotemporal points. We call these the “raw" correlations. Second, starting from the second oldest time slice, we took each of the values of a particular variable of a time slice and subtracted from them the values of the same variable at the same location but from the immediately previous time slice. We did this for all variables and then computed their pairwise correlations, which we call the correlations in differences. Finally, we took each of the values of a particular variable of a time slice and subtracted from them the values of the same variable at the same location but from the last (present-day) time slice. We then computed pairwise correlation between the resulting values for each of the variables, excluding the last time slice from the analysis (as it would just contain zeroes). We call these the correlations in “anomalies", in the sense that the resulting values represent anomalies of a variable with respect to its present-day value at a given location.

### Spatiotemporal Bayesian modeling of vegetation anomalies

We used two types of hierarchical spatio-temporal Bayesian model implemented in the R library spTimer (Bakar, Sahu, et al., 2015), in order to jointly model climate and kriged ancestry anomalies as explanatory variables for vegetation type anomalies. To simplify notation, we will now index time with the variable *t* and assume all sites have observations at the same time slices, i.e. *T*_*i*_ = *T* for all sites *s*_*i*_. We will suppose we have *n* sites, and so *nT* is the total number of spatiotemporal observations. The first of the spatiotemporal models treats the response variable ***Z***(*t*) - in our case, containing all vegetation type values in our spatial map at time *t* - as a noisy observation of a Gaussian process ***O***(*t*):

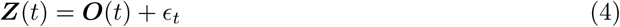

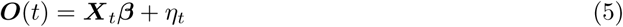

Here, ***β*** is a *p* × 1 vector of coefficients, ***X***_***t***_ is a *n* × *p* matrix of covariates at time *t*, *ϵ*_*t*_ is an error vector that only depends on an unknown pure error variance *σ*_*ϵ*_:

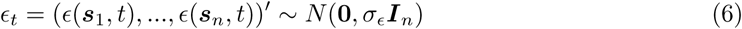

while *η*_*t*_ is a spatiotemporal nugget vector that is independent of *ϵ*_*t*_ and whose distribution depends on a site invariant spatial variance *σ*_*η*_ and the spatial correlation matrix *S*_*η*_:

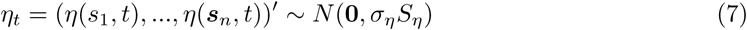

The correlation matrix *S*_*η*_ is obtained from the general Matérn correlation function (Matérn, 1986), whose shape depends on two unknown parameters - *λ* and *ν*. These control the rate of decay of the correlation as the distance between sites increases and the smoothness of the random field, respectively (Bakar, Sahu, et al., 2015).

The second model is a temporal auto-regressive model that works by incorporating a term in equation 7 that depends on the previous instance of the ***O***() process and a temporal correlation parameter *ρ*:

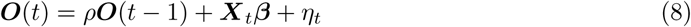

spTimer can fit these models via Gibbs sampling and infer the posterior distribution of the unknown parameters ***β***, *ϵ*_*t*_, *η*_*t*_, *ν*, *φ* and *ρ*. Unless otherwise stated, we used spTimer’s default prior distributions for these parameters, which are detailed in Bakar, Sahu, et al. (2015). Before inputting all explanatory and response variables into either model, we first centered and scaled them to have mean 0 and variance 1. We tried three different types of prior distributions for the spatial decay parameter of the Matérn correlation function (Figures S11) - a fixed value, a Uniform distribution and a Gamma distribution, each with default hyperparameters - and compared their performance using the root mean squared error of the predictions (see below).

### Assessment of prediction error

We randomly removed 20% of the grid points in the map and fitted a spatio-temporal model to the remainer of the data. We computed the root mean square error by comparing the predicted values across all temporal slices with the previously removed observed values. We then selected the spatiotemporal model (AR vs GP) and the prior distribution for the spatial decay parameter of the Matérn correlation function (fixed value vs. Uniform vs. Gamma) based on visual comparison of the root mean squared error plots for each of these model choices (Figure S11) (Bakar, Sahu, et al., 2015).

### Nonparametric bootstrapping of parameter estimates

The Gibbs sampler allows us to obtain posterior estimates of the ***β*** parameters relating the explanatory to the response variables. However, it relies on the kriged ancestry grid-point maps as input, so it does not account for the uncertainty in the estimation of these maps from the ancient genomes that we currently have. To address this, we derived confidence intervals on the Bayesian posterior estimates using a nonparametric bootstrapping approach. We created 100 pseudo-samples, by randomly sampling ancient and present-day genomes 100 times - with replacement - from among the list of all ancient and present-day genomes, then obtaining their ancestry assignments and kriging them on the spatiotemporal grid. We then fitted the Bayesian spatiotemporal model to each pseudosample, and thus obtained a distribution of bootstrapped ***β*** parameter estimates, from which we obtained 95% confidence intervals.

### Arrival time maps

We created ancestry arrival time maps by recording the time in each cell of the spatial grid at which the spatiotemporal surface map reaches a kriged ancestry value higher than X, where X could be 50% or 75%. In this case, we used a spatial grid of 5,000 points and 200-year time intervals. To create the cultural arrival maps for the spread of farming, we overlaid a 50×50km map covering Europe and selected, for each square, the oldest radiocarbon date directly associated with early farming. The dataset we used to obtain these dates came from the EUROFARM database, which contains 1,779 records of archaeological farming sites (Vander Linden in prep.). It was then spatially kriged using the spatstat package in R.

### Front speed estimation

To estimate the front speed of the spread of NEOL and YAM ancestries, we used a method developed by Pinhasi, Fort, and Ammerman (2005) and Silva and Steele (2014), in which the authors regress great-circle distances of sampled locations to a hypothesized migration origin against the time at which the migration reached those locations. The negative inverse of the slope is then an estimate of the migration front speed. We restricted to genomes older than 5,000 years BP for the NEOL ancestry spread and to genomes older than 3,000 years BP for the YAM ancestry spread. We used Cayönü (37.38N, 40.39E) as the NEOL ancestry origin, based on estimates of the Neolithic farmer expansion origin (Pinhasi, Fort, and Ammerman, 2005; Silva and Steele, 2014). We set various points at the center and extremes of the hypothesized original Yamnaya distribution in the Eurasian steppe as the YAM ancestry origin (Table 1). We used a ranged major axis (RMA) regression approach implemented in the R package lmodel2 (Borcard, Gillet, and Legendre, 2018), which assumes a symmetrical distribution of measurement error in both distance and time.

## Supporting information

Supplementary Animation 3

Supplementary Animation 2

Supplementary Animation 1

## Acknowledgments

We thank John Novembre, Rasmus Nielsen, Michael K. Borregaard, Mark G. Thomas and Kurt H. Kjær for helpful advice and discussions. FR was funded by a Villum Young Investigator award (project no. 000253000). The palaeo-vegetation research was funded by the Leverhulme Trust and we gratefully acknowledge contributors to the European Pollen Database.

---

**Figure S1:**
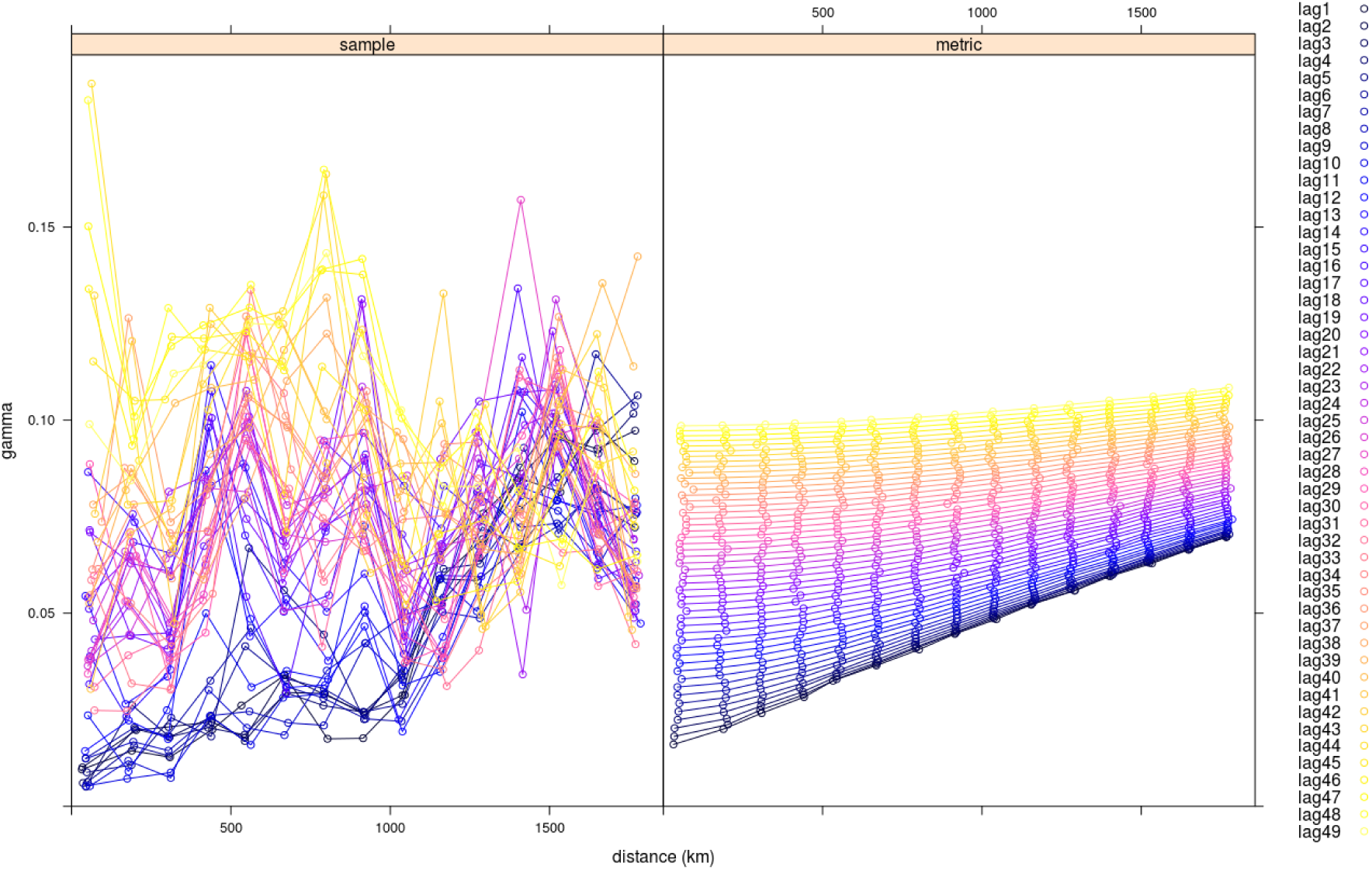
Spatiotemporal semivariogram of YAM ancestry in Europe, with temporal lags of 60-year increments. Left panel: empirical semivariogram. Right panel: fitted semivariogram model using the metric method. The lags are in 60-year increments.

**Figure S2:**
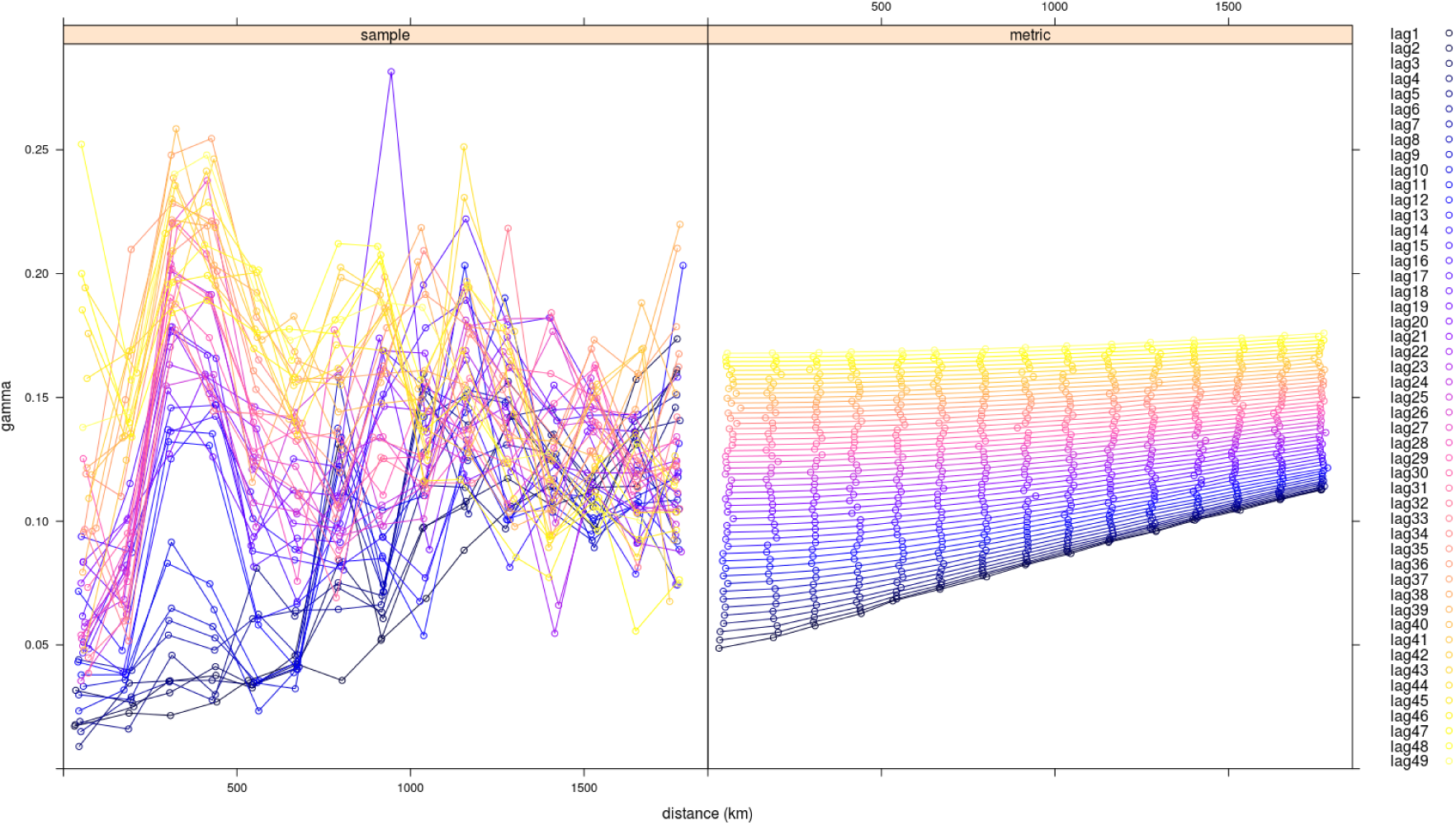
Spatiotemporal semivariogram of NEOL ancestry in Europe, with temporal lags of 60-year increments. Left panel: empirical semivariogram. Right panel: fitted semivariogram model using the metric method. The lags are in 60-year increments.

**Figure S3:**
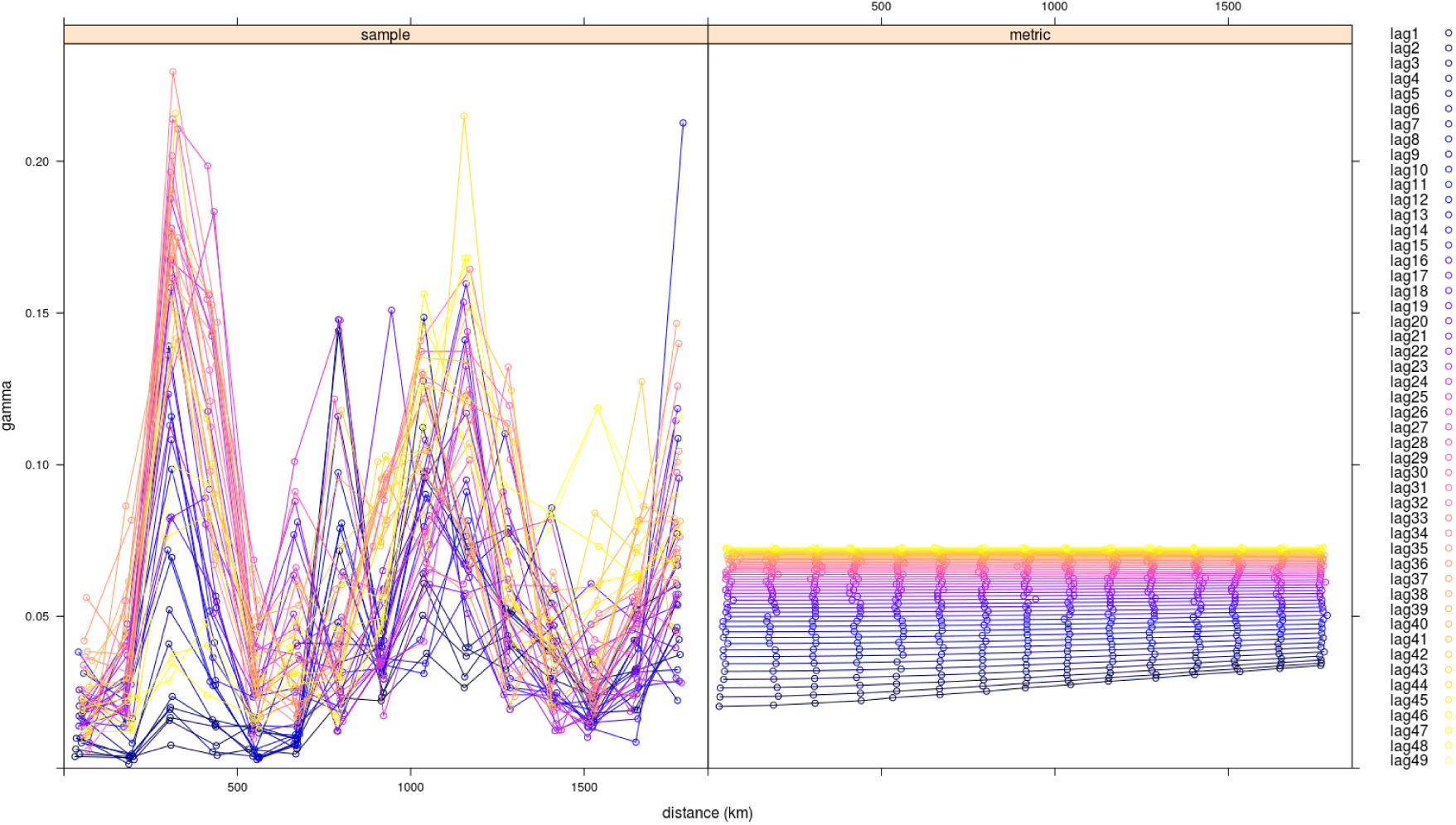
Spatiotemporal semivariogram of HG ancestry in Europe, with temporal lags of 60-year increments. Left panel: empirical semivariogram. Right panel: fitted semivariogram model using the metric method. The lags are in 60-year increments.

**Figure S4:**
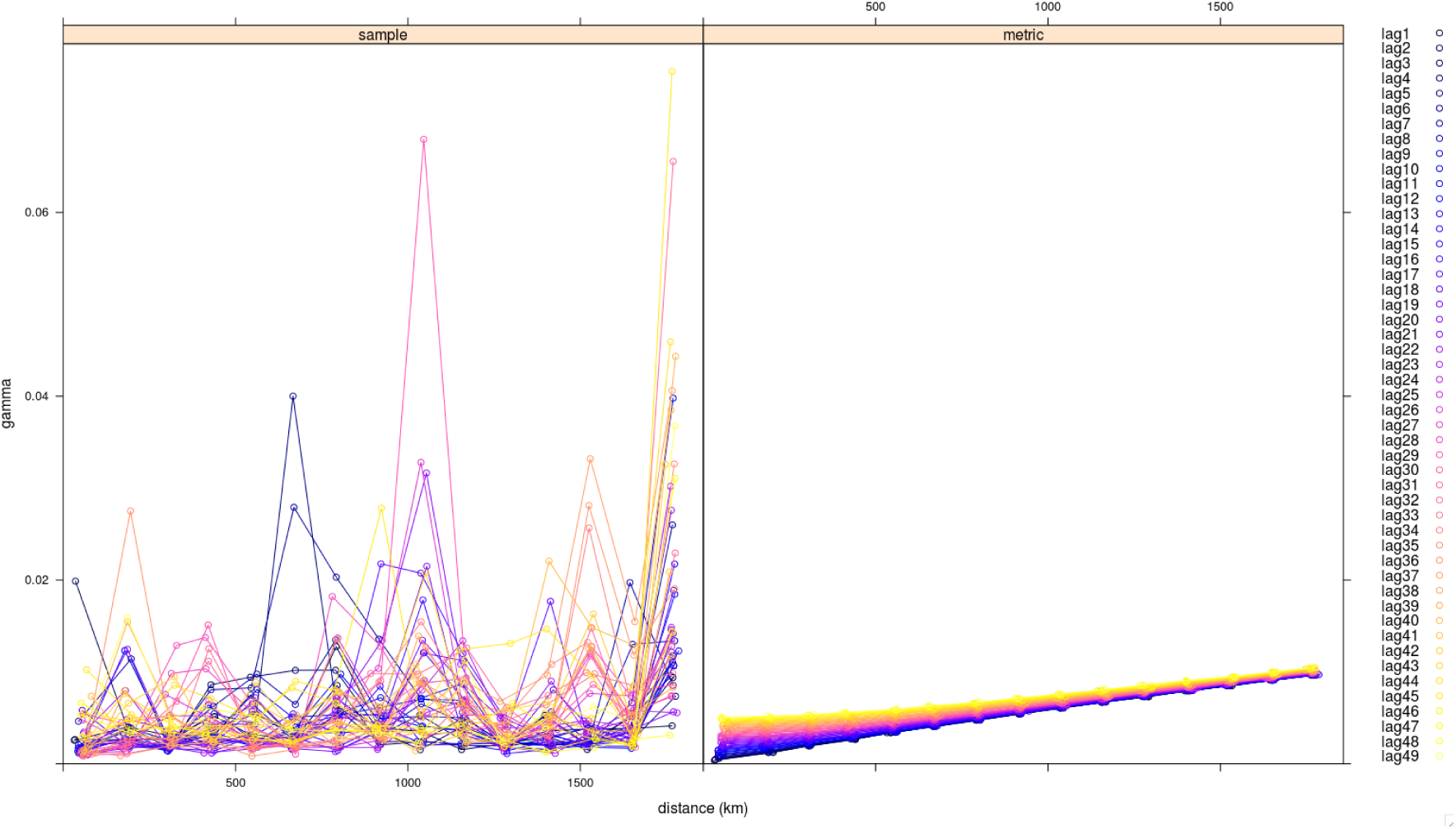
Spatiotemporal semivariogram of NAF ancestry, with temporal lags of 60-year increments. Left panel: empirical semivariogram. Right panel: fitted semivariogram model using the metric method. The lags are in 60-year increments.

**Figure S5:**
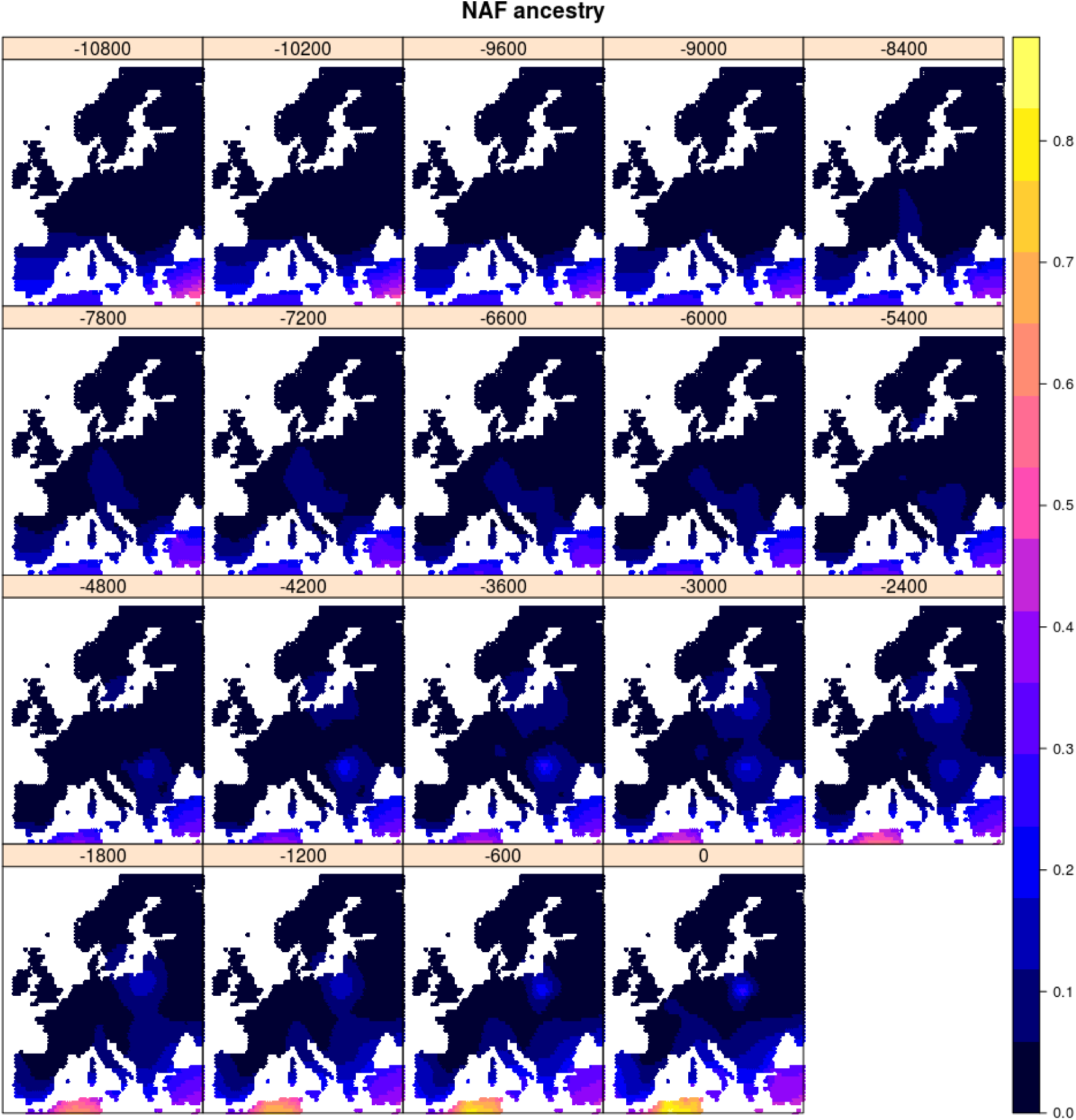
Spatiotemporal kriging of NAF ancestry during the Holocene, using 5000 spatial grid points. The colors represent the predicted ancestry proportion at each point in the grid.

**Figure S6:**
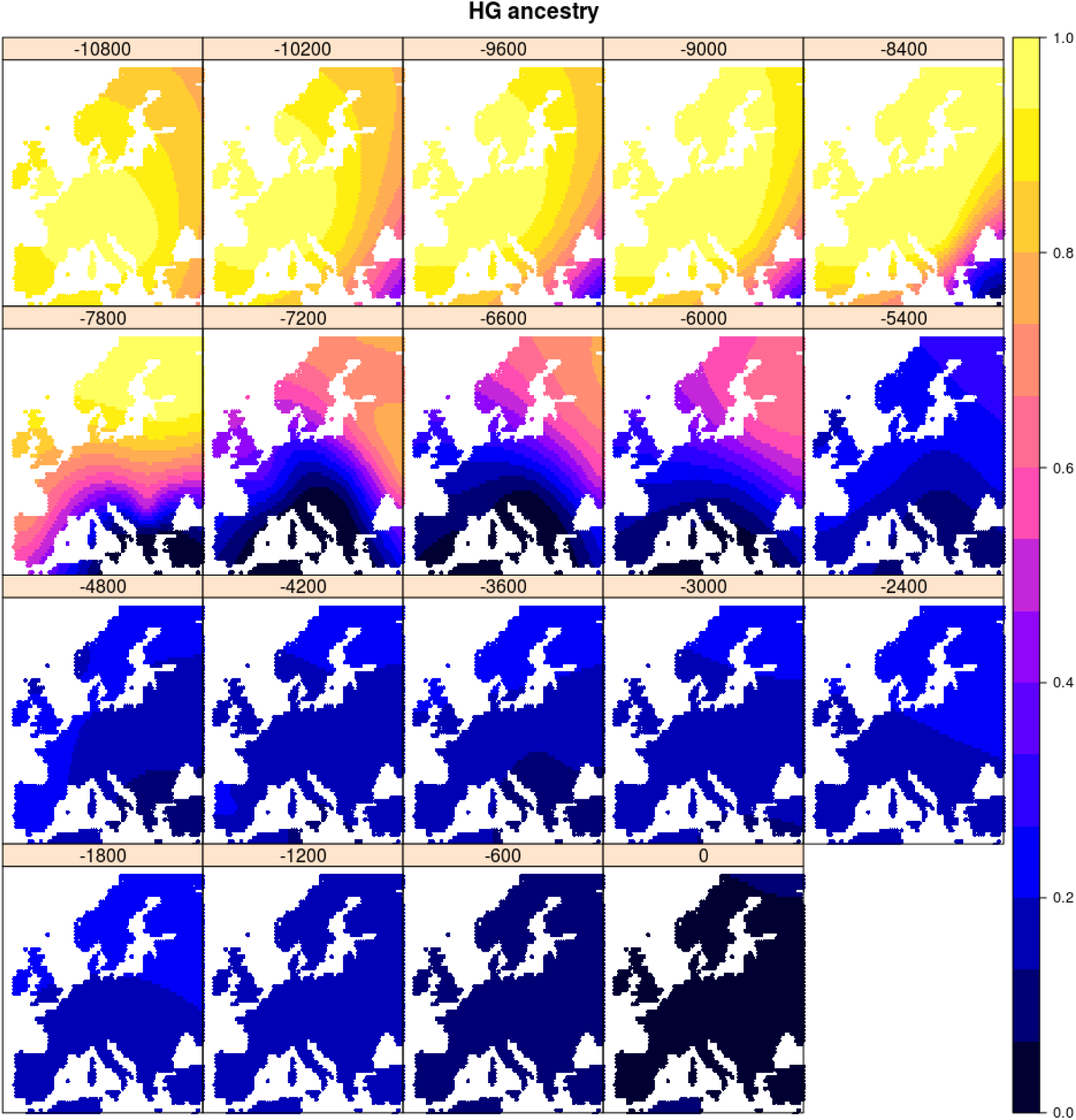
Spatiotemporal kriging of HG ancestry during the Holocene, using 5000 spatial grid points. The colors represent the predicted ancestry proportion at each point in the grid.

**Figure S7:**
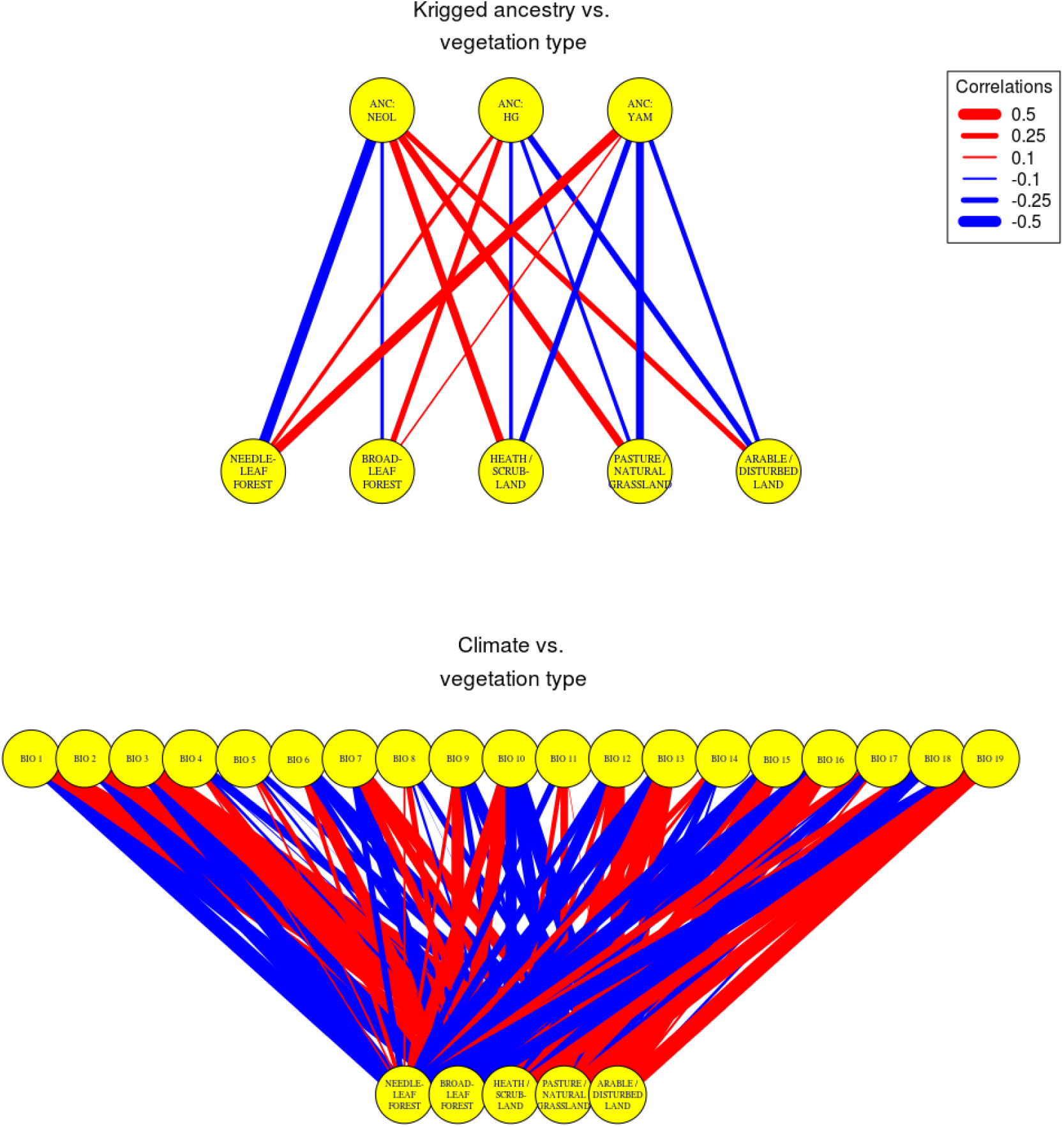
Correlations between kriged ancestry and vegetation type (top) or between climate variables and vegetation type (bottom), computed across time and space. ANC = ancestry. NEOL = neolithic farmer. HG = hunter-gatherer. YAM = Yamnaya. The climate variables follow the WorldClim nomenclature. BIO1 = Annual Mean Temperature. BIO2 = Mean Diurnal Range (Mean of monthly (max temp - min temp)). BIO3 = Isothermality (BIO2/BIO7) (* 100). BIO4 = Temperature Seasonality (standard deviation *100). BIO5 = Max Temperature of Warmest Month. BIO6 = Min Temperature of Coldest Month. BIO7 = Temperature Annual Range (BIO5-BIO6). BIO8 = Mean Temperature of Wettest Quarter. BIO9 = Mean Temperature of Driest Quarter. BIO10 = Mean Temperature of Warmest Quarter. BIO11 = Mean Temperature of Coldest Quarter. BIO12 = Annual Precipitation. BIO13 = Precipitation of Wettest Month. BIO14 = Precipitation of Driest Month. BIO15 = Precipitation Seasonality (Coefficient of Variation). BIO16 = Precipitation of Wettest Quarter. BIO17 = Precipitation of Driest Quarter. BIO18 = Precipitation of Warmest Quarter. BIO19 = Precipitation of Coldest Quarter.

**Figure S8:**
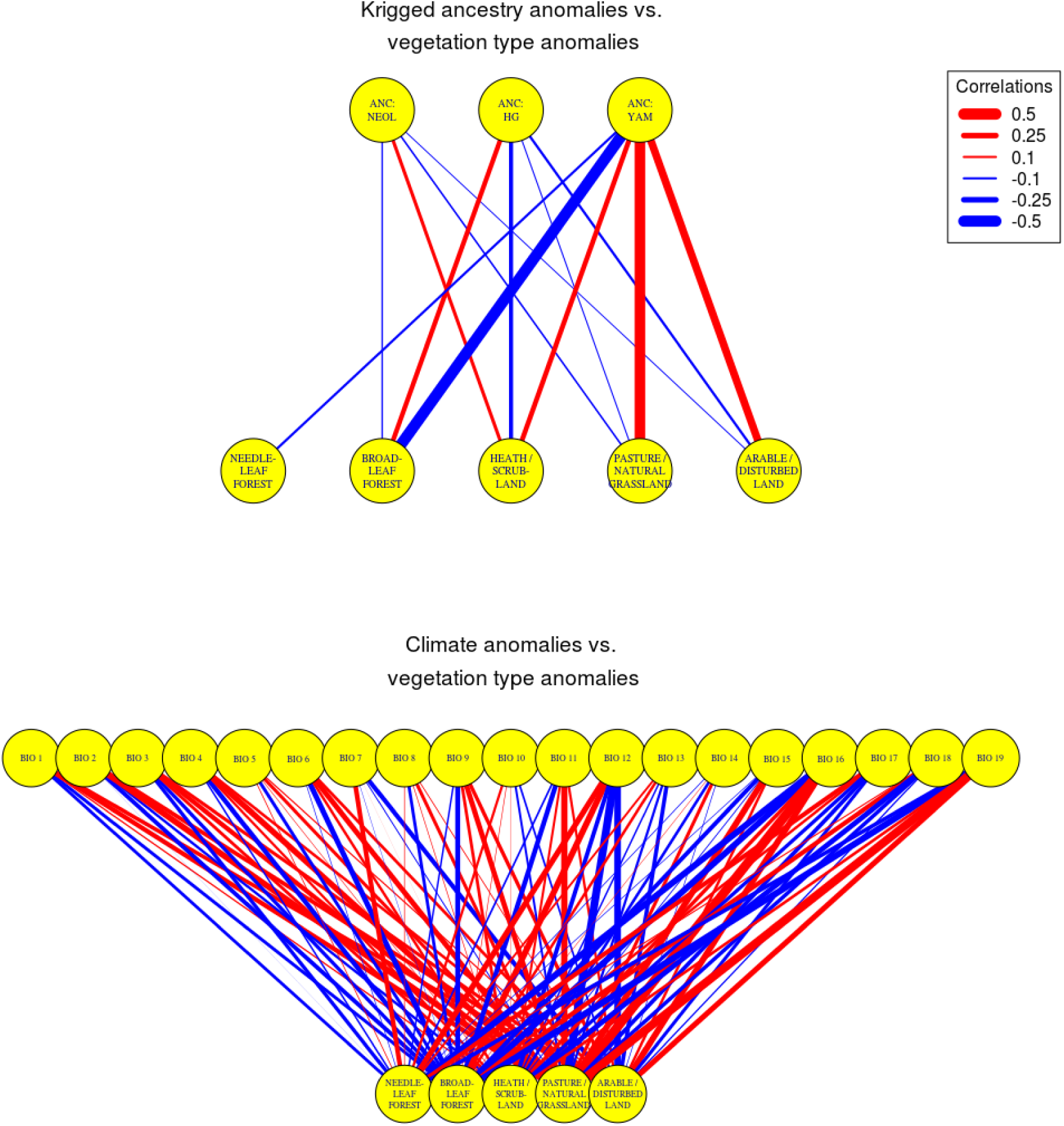
Correlations between anomalies, defined as the values of a variable in each past time slice before the present, after subtracting the value of that variable at the same location in the present. Top: Correlations between kriged ancestry anomalies and vegetation type anomalies. Bottom: Correlations between climate anomalies and vegetation type anomalies. ANC = ancestry. NEOL = neolithic farmer. HG = hunter-gatherer. YAM = Yamnaya. The climate variables follow the WorldClim nomenclature. BIO1 = Annual Mean Temperature. BIO2 = Mean Diurnal Range (Mean of monthly (max temp - min temp)). BIO3 = Isothermality (BIO2/BIO7) (* 100). BIO4 = Temperature Seasonality (standard deviation *100). BIO5 = Max Temperature of Warmest Month. BIO6 = Min Temperature of Coldest Month. BIO7 = Temperature Annual Range (BIO5-BIO6). BIO8 = Mean Temperature of Wettest Quarter. BIO9 = Mean Temperature of Driest Quarter. BIO10 = Mean Temperature of Warmest Quarter. BIO11 = Mean Temperature of Coldest Quarter. BIO12 = Annual Precipitation. BIO13 = Precipitation of Wettest Month. BIO14 = Precipitation of Driest Month. BIO15 = Precipitation Seasonality (Coefficient of Variation). BIO16 = Precipitation of Wettest Quarter. BIO17 = Precipitation of Driest Quarter. BIO18 = Precipitation of Warmest Quarter. BIO19 = Precipitation of Coldest Quarter.

**Figure S9:**
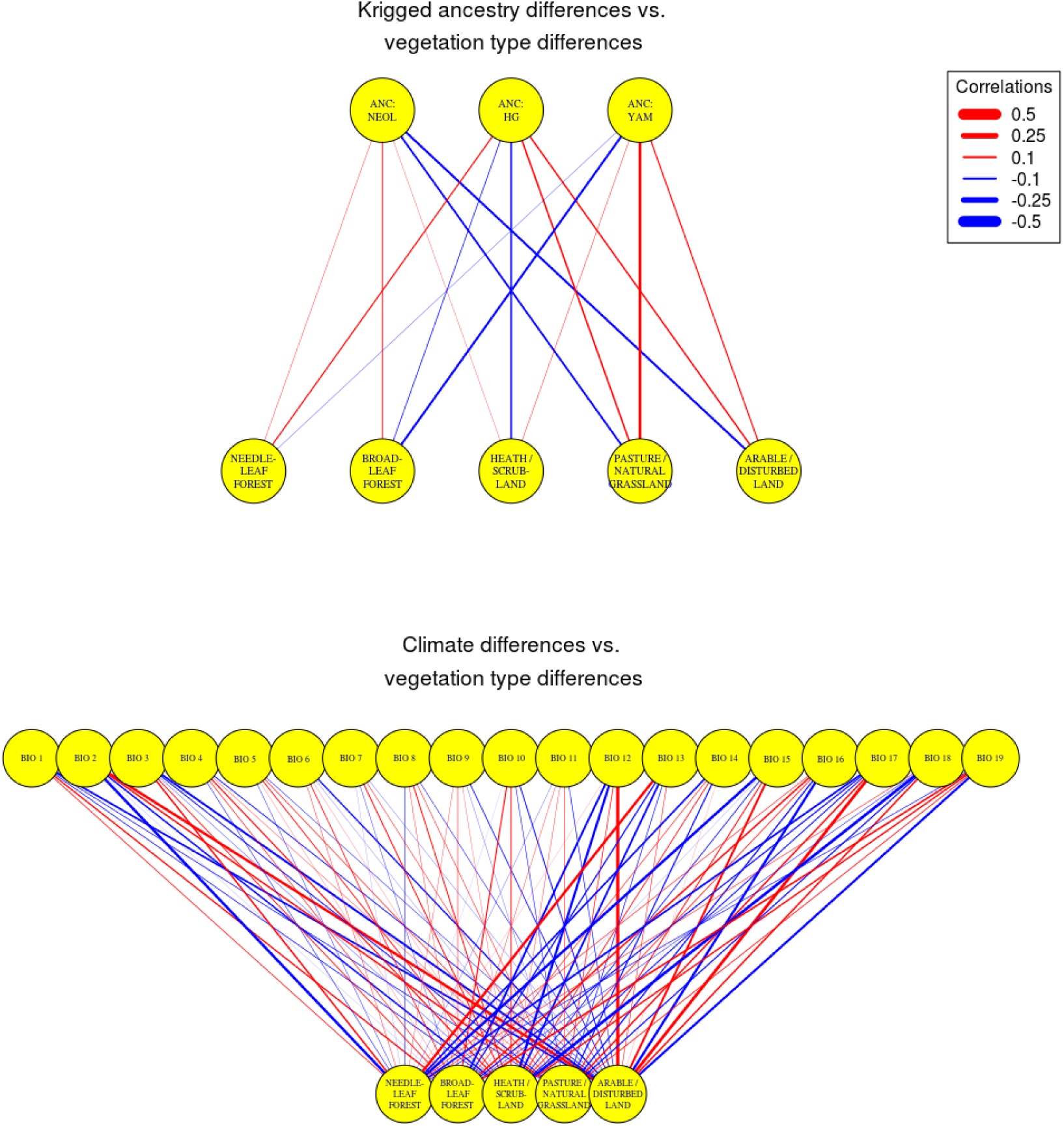
Correlations between differences in kriged ancestry and vegetation type at each time slice (top) or between differences in climate variables and in vegetation type at each time slice (bottom), computed across time and space. ANC = ancestry. ANC:NEOL = neolithic farmer ancestry. ANC:HG = hunter-gatherer ancestry. ANC:YAM = Yamnaya steppe ancestry. The climate variables follow the WorldClim nomenclature. BIO1 = Annual Mean Temperature. BIO2 = Mean Diurnal Range (Mean of monthly (max temp - min temp)). BIO3 = Isothermality (BIO2/BIO7) (* 100). BIO4 = Temperature Seasonality (standard deviation *100). BIO5 = Max Temperature of Warmest Month. BIO6 = Min Temperature of Coldest Month. BIO7 = Temperature Annual Range (BIO5-BIO6). BIO8 = Mean Temperature of Wettest Quarter. BIO9 = Mean Temperature of Driest Quarter. BIO10 = Mean Temperature of Warmest Quarter. BIO11 = Mean Temperature of Coldest Quarter. BIO12 = Annual Precipitation. BIO13 = Precipitation of Wettest Month. BIO14 = Precipitation of Driest Month. BIO15 = Precipitation Seasonality (Coefficient of Variation). BIO16 = Precipitation of Wettest Quarter. BIO17 = Precipitation of Driest Quarter. BIO18 = Precipitation of Warmest Quarter. BIO19 = Precipitation of Coldest Quarter.

**Figure S10:**
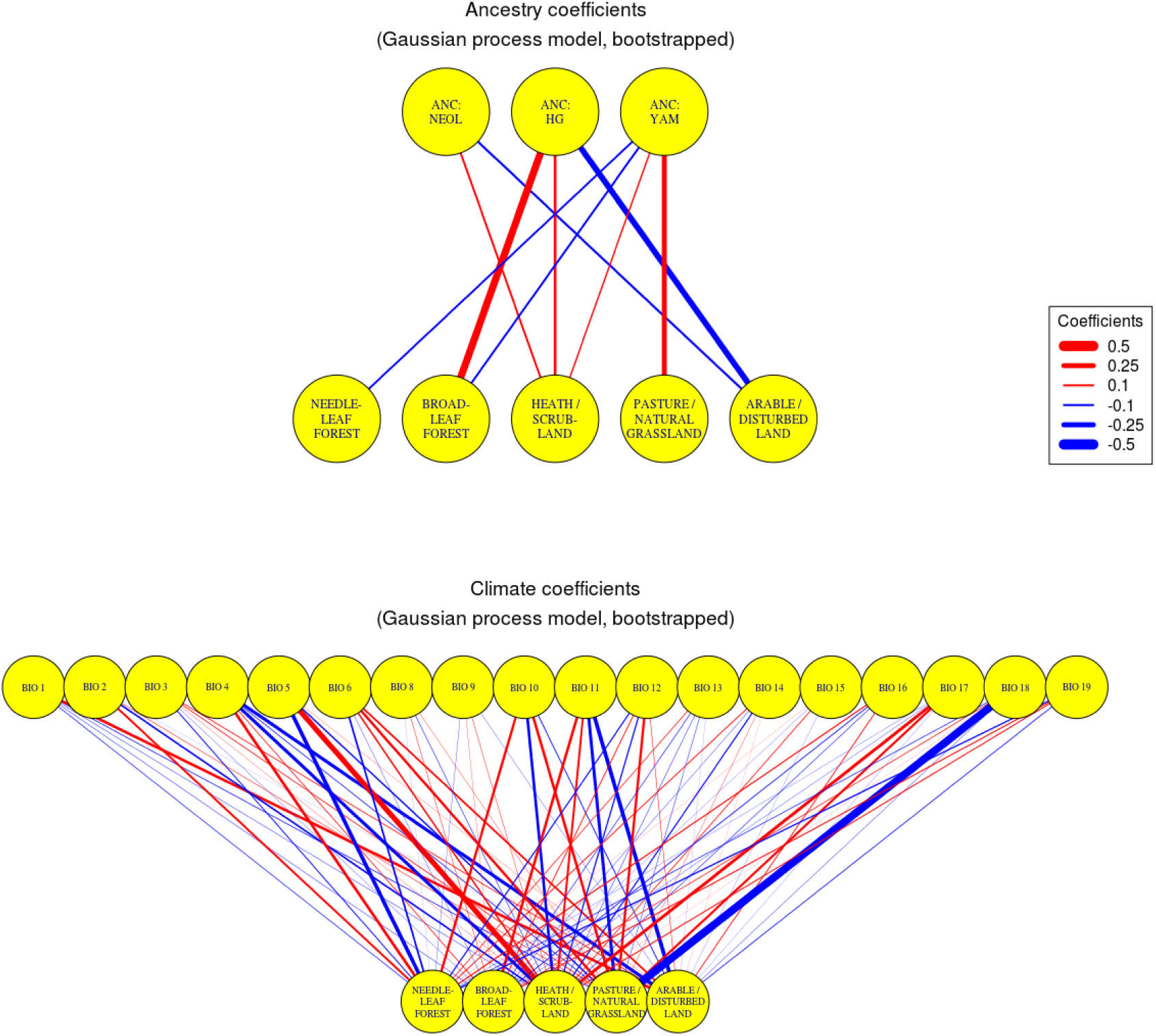
Spatiotemporal Gaussian process model for paleo-vegetation anomalies, using kriged ancestry anomalies and anomalies from simulation-based paleo-climate reconstructions as explanatory variables. Here, we used bootstrapping to account for uncertainty in the ancestry kriging estimation. The depicted lines denote the mean coefficients from the bootstrap distribution. Coefficients whose bootstrap distribution has a 95% central probability mass interval that spans the value of 0 are not depicted. ANC = ancestry. NEOL = neolithic farmer. HG = hunter-gatherer. YAM = Yamnaya. CLIM = Climate. The climate variables follow the WorldClim nomenclature.

**Figure S11:**
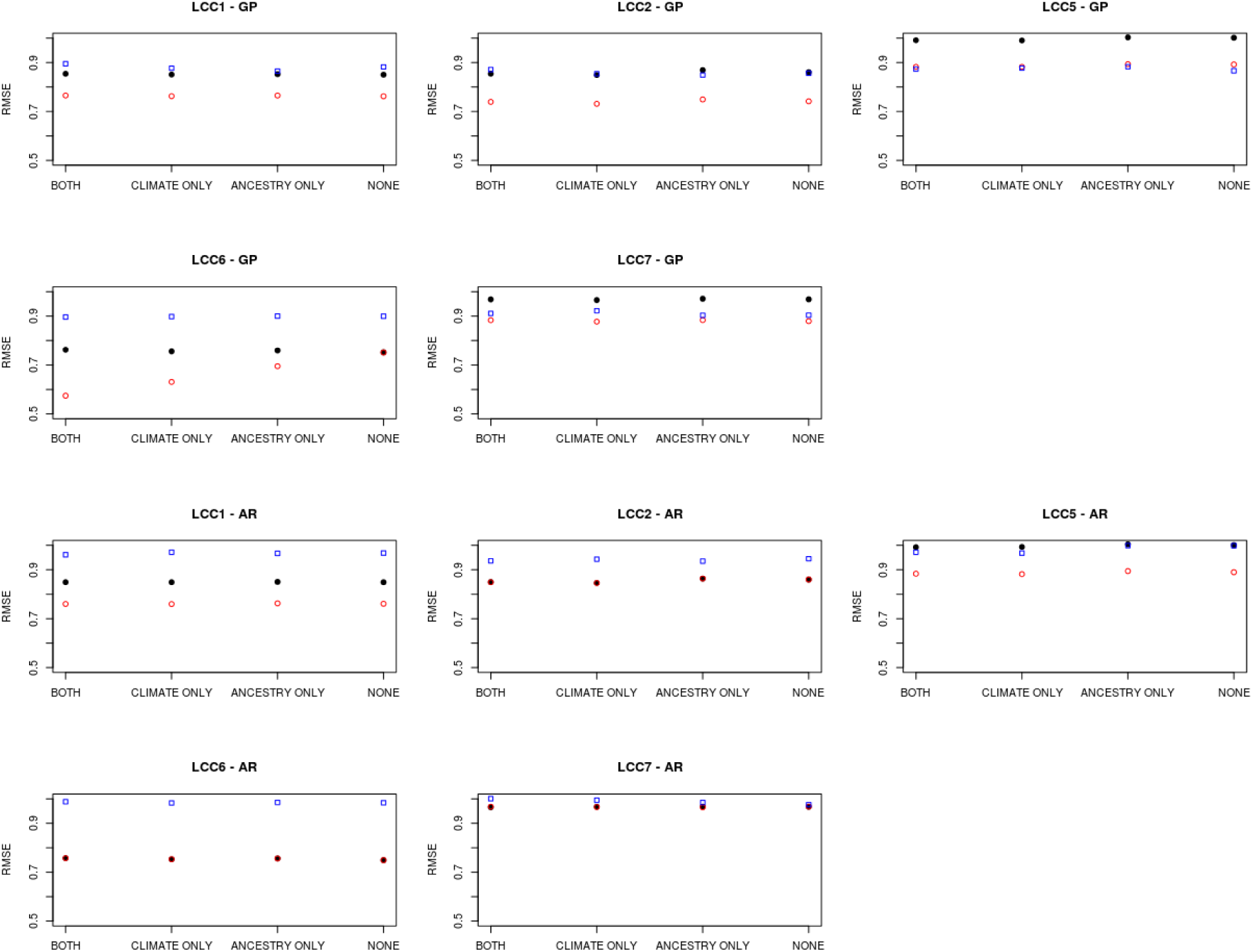
Root mean squared error for each fitted spatio-temporal model for paleo-vegetation anomalies. Black filled circles: using a fixed spatial decay parameter of the Matérn correlation function (default value in spTimer). Red circles: using a Uniform prior distribution for the spatial decay parameter (with default hyperparameters in spTimer). Blue squares: using a Gamma prior distribution for the spatial decay parameter (with default hyperparameters in spTimer). GP = Gaussian process model. AR = autoregressive model. BOTH = model incorporating both climate and ancestry explanatory variables. CLIMATE ONLY = model incorporating climate explanatory variables. CLIMATE ONLY = model incorporating ancestry explanatory variables. NONE = model incorporating neither ancestry nor climate explanatory variables.

