## Supplementary figures and images for "A geostatistical approach to modelling human Holocene migrations in Europe using ancient DNA"

### Supplementary Animation 1

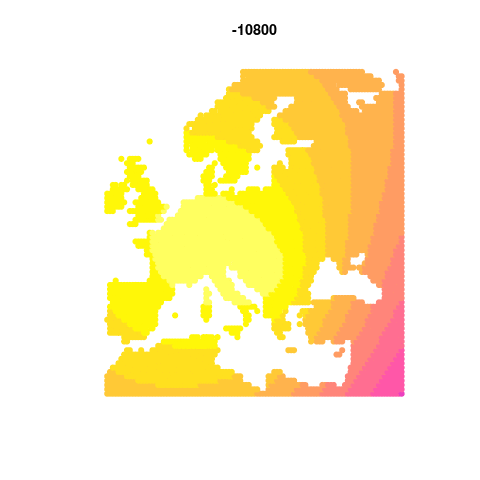

### Supplementary Animation 2

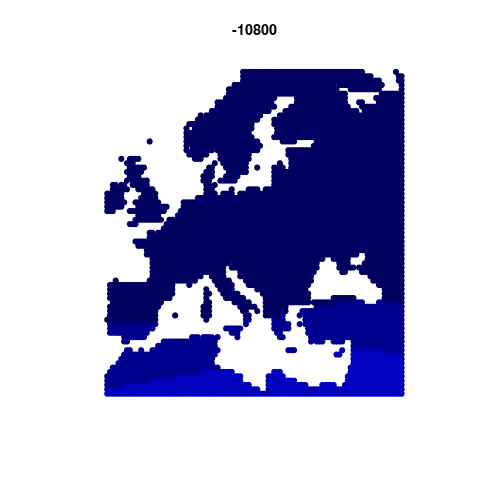

### Supplementary Animation 3

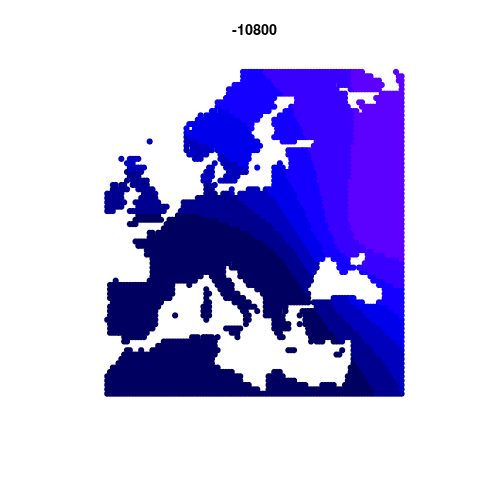
